# A bHLH interaction code controls bipotential differentiation and self-renewal in the *Drosophila* gut

**DOI:** 10.1101/685347

**Authors:** Aleix Puig-Barbe, Svenja Dettmann, Vinicius Dias Nirello, Helen Moor, Sina Azami, Bruce A. Edgar, Patrick Varga-Weisz, Jerome Korzelius, Joaquín de Navascués

## Abstract

Multipotent adult stem cells balance self-renewal with differentiation into various cell types. How this balance is regulated at the transcriptional level is poorly understood. Here we show that a network of basic Helix-Loop-Helix (bHLH) transcription factors controls both stemness and bi-potential differentiation in the *Drosophila* adult intestine. We find that homodimers of Daughterless (Da), homolog of mammalian E proteins, maintain self-renewal of intestinal stem cells (ISCs), antagonising the Enteroendocrine fate promoted by heterodimers of Da and Scute (Sc, homolog of ASCL). The HLH factor Extramacrochaetae (Emc, homologous to Id proteins) promotes absorptive differentiation by titrating Da and Sc. Emc prevents the committed absorptive progenitor from de-differentiating, underscoring the plasticity of these cells. Switching physical interaction partners in this way enables the active maintenance of stemness while priming stem cells for differentiation along two alternative fates. Such regulatory logic is likely operative in other bipotent stem cell systems.

## INTRODUCTION

The regulation of stem cell fate decisions hinges on transcriptional control by sequence-specific transcription factors (TFs) forming gene regulatory networks that steer cells along particular differentiation trajectories (Graf and Enver, 2009; Levine and Davidson, 2005; Moris et al., 2016). These trajectories are often considered a succession of binary steps regulated by cross-antagonism between TF pairs (Graf and Enver, 2009; Simon et al., 2018). However, active multipotent stem cells need to decide between the maintenance of their stem identity and several options of commitment into distinct mature cell fates. To understand how multipotent stem cells make these choices, knowledge of the functional interactions between transcription factors is essential (Cinquin and Demongeot, 2005; 2002).

Intestinal stem cells (ISCs) are a paradigm of multipotency in adult tissues. ISCs face a choice between self-renewal and differentiation into either the secretory or absorptive cell lineage (reviewed in Crosnier et al., 2006; Jiang and Edgar, 2012; Philpott and Winton, 2014). The intestinal secretory lineage in *Drosophila* consists of enteroendocrine cells (EE) (Micchelli and Perrimon, 2006; Ohlstein and Spradling, 2006). Absorptive cells are called enterocytes (ECs) and differ in morphology and function along the anterior-posterior axis of the gut (Buchon et al., 2013; Marianes and Spradling, 2013). In *Drosophila*, ISCs produce lineage-specific precursors through distinct molecular triggers. High Notch signalling induces formation of enteroblasts (EBs), which will give rise to ECs. Expression in ISCs of the bHLH (basic Helix-Loop-Helix) transcription factors Scute (Sc) or Asense (Ase), members of the *achaete-scute Complex* (*AS-C*, homologs of *ASCL* mammalian genes) induces the formation of EE precursors (pre-EEs), which quickly turn into EE cells (Bardin et al., 2010; Chen et al., 2018; Zeng and Hou, 2015). It is not clear whether upon ISC division, daughter cells first commit to differentiation before choosing a lineage (two consecutive binary decisions), or are already lineage-primed before they lose self-renewing ability (binary decisions in inverse order), or choose at the same time between self-renewal and two potential lineages to commit to (single triple decision) (Gervais and Bardin, 2017; Perdigoto et al., 2011). Little is known about the molecular mechanisms that could allow a triple decision between self-renewal and bi-potential differentiation.

The bHLH family of transcription factors control cell fate in multiple developmental contexts (reviewed in Bertrand et al., 2002; Murre, 2005; Murre et al., 1994). Their HLH motif mediates dimerization, while the preceding region, rich in basic amino acids, allows DNA binding (reviewed in Massari and Murre, 2000). Class I of bHLH factors comprises proteins such as E47, E2-2 and HEB (E proteins, encoded by *TCF3/4/12* in mammals; *daughterless* —*da*— in *Drosophila*). Class I bHLH proteins can make dimers within their class (e.g., Da:Da, or E47:HEB), but also heterodimerise with class II bHLH factors. By contrast, class II bHLH factors such as MYOD or ASCL (MyoD and proteins encoded in the *AS-C* in *Drosophila*) usually form trans-activating complexes only in heterodimers with class I factors (Murre et al., 1994). Both class I and class II factors can heterodimerise with class V proteins (reviewed in Campuzano, 2001; Wang and Baker, 2015). The mammalian members of this class are named *Inhibitors of DNA binding* (Id proteins) because they lack the stretch of basic amino acids preceding the HLH domain, rendering their heterodimers unable to bind DNA (Murre et al., 1994). Their only homolog in *Drosophila* is *extra macrochaetae* (*emc*).

Class II bHLH factors regulate differentiation in the metazoan intestine (reviewed in Hartenstein et al., 2017; Philpott and Winton, 2014). In *Drosophila*, Sc and Ase can initiate EE differentiation (Amcheslavsky et al., 2014; Bardin et al., 2010; Chen et al., 2018; Z. Guo and Ohlstein, 2015), while other bHLH factors maintain EE function (Dimmed, homolog of NeuroD; Beebe et al., 2015), or promote their functional diversity (Tap, homolog of Neurogenin-3; Hartenstein et al., 2017). On the other hand, Da is required for ISC maintenance, as ISCs mutant for *da* differentiate (Bardin et al., 2010; Lan et al., 2018). However, the interaction partners of Da to maintain stemness are not known, and how different bHLH factors dimerise to control differentiation has not been explored. Here we identify the Da homodimer as the critical bHLH complex maintaining ISC self-renewal and find a role of Emc in titrating Da and Sc to promote absorptive differentiation. We show that Da:Da and Da:Sc dimers functionally cooperate promoting ISC fate, but act antagonistically for EE differentiation. Our results reveal a network of bHLH factors that forms a three-way switch to regulate self-renewal and bipotential differentiation in the adult fly gut.

## RESULTS

### Da homodimers maintain stemness and prevent differentiation

We quantified the effect of *da* on differentiation in individual null *da^10^* clones using the MARCM method (**Figure S1A-B**; Lee and Luo, 1999) and in the entire ISC/EB population by RNAi-mediated knock-down of *da* followed by lineage tracing using the *escargot* Flip-Out approach (“*esg^TS^-FO*”, see **Figure S1C**; Jiang et al., 2009). Loss of *da* led to almost complete differentiation of cells into ECs, and occasionally into EEs, in *da^10^* mutant clones (**Figure 1A-B, I** and **Tables S1-2**) and in the *esg^+^* cells expressing *da^RNAi^*(**Figure 1C-E, I**; cell type markers as in **Figure S1D**). By contrast, in ISCs and EBs that overexpressed *da* with *esg^TS^-FO*, differentiation into ECs was almost completely impaired, while EE differentiation increased (**Figure 1F, I**). Importantly, only a few *esg^+^* cell nests overexpressing *da* survived and this was rescued by co-expression of apoptosis inhibitor p35 (**Figure 1J-K**). While the cell composition of *esg^TS^-FO > da* tissue was very different to the wild type, the size of GFP^+^ clusters expressing *da* was like in the controls: most remained below 5 cells, and a few became considerably larger (**Figure 1L**).

**Figure 1.**
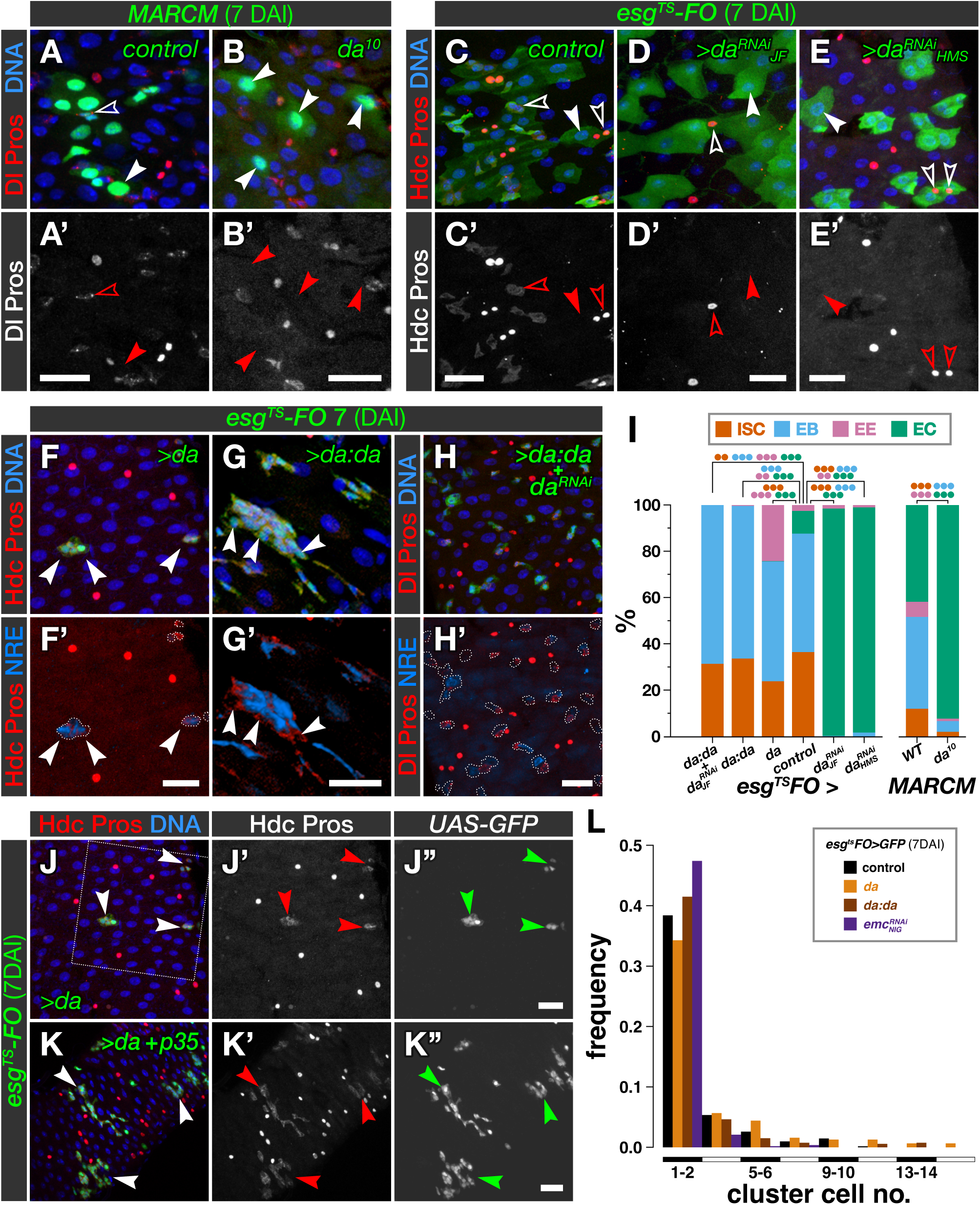
Da homodimers are required and sufficient to maintain ISCs undifferentiated. **A-B.** Cells in MARCM clones for *da^10^* (B) are mostly ECs. ISCs, EBs and EEs are dramatically reduced respect to wild-type clones (A). Panel B is reproduced in Figure 6A to aid comparison. **C-E.** Expressing *da^RNAi^* with *esg^TS^-FO* using transgenes *JF02488* (D) and *HMS01851* (E) leads to the differentiation of most cells into ECs, occasionally into EEs. Controls (C) maintain ISCs and EBs. Solid and empty arrowheads (A-E): ECs and ISCs or EEs, respectively. **F-G.** Overexpression of *da* (F) or *da:da* (G) with *esg^TS^-FO* prevents formation of ECs; *da* overexpression allows EE differentiation and strongly reduces *esg^+^* cell nest density. Solid arrowheads (F-G): ISCs/EBs. **H.** Overexpression of *da:da* with *esg^TS^-FO* while depleting endogenous Da with *JF02488* interferes blocks all differentiation. **I.** Cell composition of GFP^+^ tissue and clones from A-H. Some data are reproduced in Figure 7I-J to aid comparison. **J-K.** Expression of *da* with *esg^TS^-FO* (J) results in *esg^+^* cell death, with few survivors that cannot differentiate; co-expression of *p35* (K) rescues cell death, but not differentiation. J is a wider field of view than the tissue shown in panel F. Arrows: *esg^+^* Hdc^+^ ISCs/EBs. **L.** Histograms of GFP^+^ cluster sizes for *esg^TS^-FO* driving expression of *da*, *da:da*, *emc^RNAi^* and their control. The X axis is truncated at 16 cells, but there were larger clusters in all conditions except *emc^RNAi^*, at frequencies of 0.3% or lower. DAI: days after induction. Scale bars: 20µm. *p*-values (binomial regression for individual cell types): ● < 0.05; ●● < 0.01; ●●● < 0.001. See **Tables S1-2** for statistical details.

We sought to determine the identity of the Da partners involved in preventing EC differentiation. Since Da can form homodimers to control differentiation and proliferation (reviewed in Wang and Baker, 2015), we overexpressed forced Da homodimers using a *da:da* tethered construct (Tanaka-Matakatsu et al., 2014) with *esg^TS^-FO* to test their capacity to block differentiation. The resulting GFP^+^ tissue comprised mostly ISCs and EBs, which distributed in clusters of similar size to the wild type (**Figure 1G, I, L**). To test that Da homodimers were enough to maintain self-renewal without other Da-containing complexes, we expressed Da:Da while removing endogenous Da with the *da^RNAi^* transgene *P{TRiP.JF02488}*, which does not target the *da:da* construct (**Figure S1E**). This prevented differentiation entirely (all cells were either ISCs or EBs; **Figure 1H-I**). In addition, we detected no *esg^+^*cell death when the tethered *da:da* construct was overexpressed (**Figures 1G-H**, compare with **Figure 1J**). Together, these results show that Da homodimers promote stemness; additionally, Da probably participates in another complex that induces *esg*^+^ cell death.

### Da:Sc and Da:Da antagonise each other in secretory differentiation

Our results so far indicate that EE differentiation requires the transition from the transcriptional program of Da:Da to that of Da:Sc. This could occur through a ‘switch’, with Da:Sc targets being epistatic over those of Da:Da, or ‘antagonism’, whereby the relative strengths of the two programs determine cell fate. To distinguish between these alternatives, we compared the effects of overexpressing *sc* with *esg^TS^-FO* with those of co-expressing *da* and *sc*. Overexpression of *sc* alone leads to the induction of Pros-positive, Dl-negative and Dl/Pros double-positive cells (**Figures 2A and S2A**), as expected (Bardin et al., 2010; Chen et al., 2018). This was likely mediated by Da:Sc, as co-expression of *sc* and *da^RNAi^* strongly diminished the induction of all Pros-positive cells (**Figure S2A-C**). Many of these cells exhibited the mitotic marker phospho-Histone H3S10 (PH3) (**Figure S2D**) and were probably trapped in a pre-EE state (Chen et al., 2018). The co-expression of *da* and *sc* greatly reduced the number of Pros-positive, Dl-negative (EE) cells and led to an increase in all Dl-positive cells (**Figure 2B, D**), while maintaining mitotic figures (**Figure S2E, G**). Since endogenous *da* is expressed weakly (Bardin et al., 2010), the overexpression of *da* and *sc* will likely produce more Da:Sc dimers than that of *sc* alone. Therefore, the suppression of the *sc* phenotype by *da+sc* must be due to a higher Da:Da-to-Da:Sc ratio. This suggests antagonism between Da:Sc and Da:Da, and predicts that the co-expression of tethered *da:da* and *sc* would result in even less EE differentiation, as endogenous Da available for Da:Sc dimers will be limiting. Indeed, we observed a ∼2-fold increase of Dl-positive, Pros-negative cells at the expense of all Pros-positive cells, especially the Dl-negative (**Figure 2C-D**), with an increase in PH3^+^ cells (**Figure S2F-G**).

**Figure 2.**
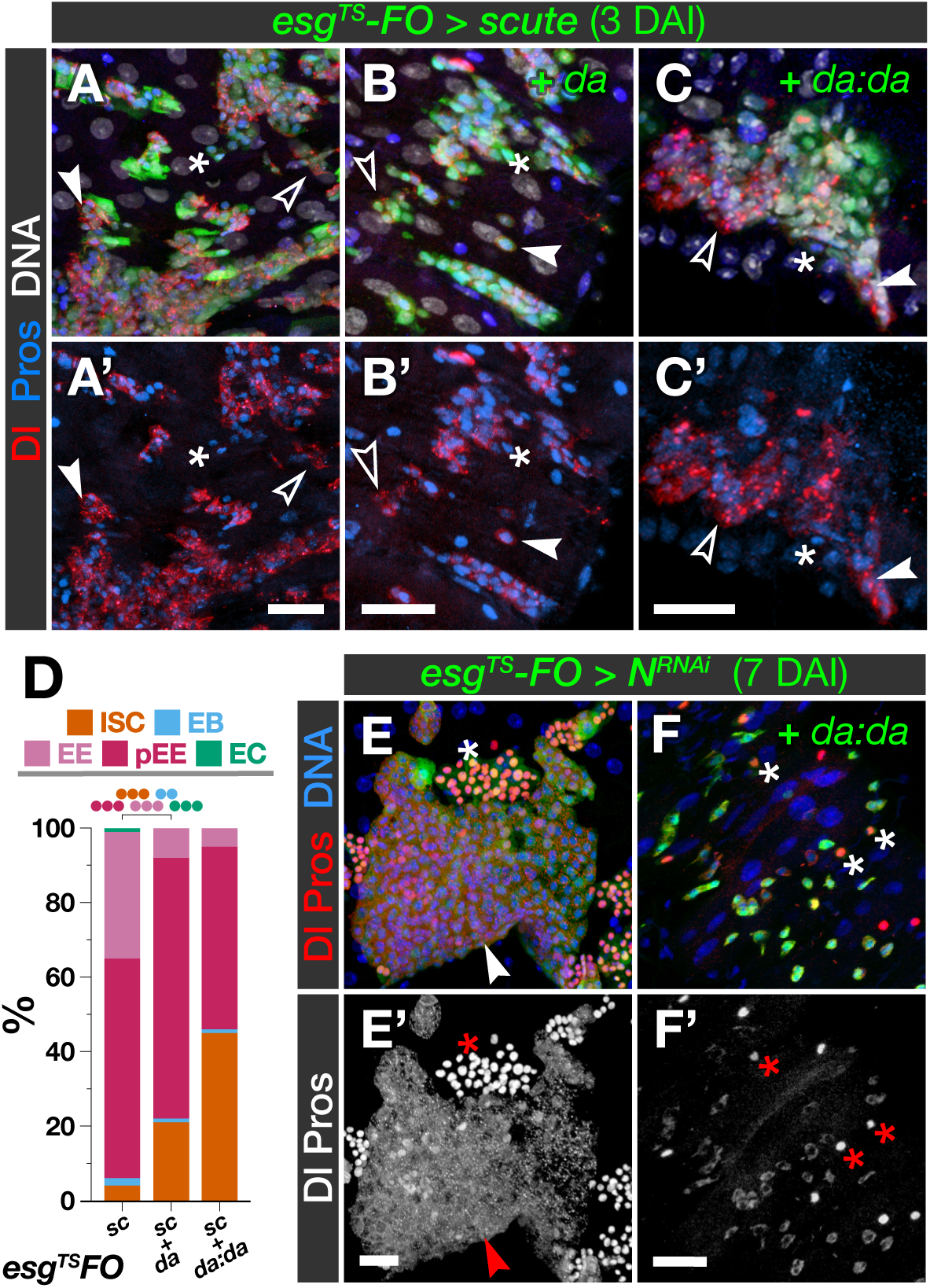
Da:Sc and Da:Da antagonise each other in EE differentiation. **A-C.** Overexpression of *sc* with *esg^TS^-FO* leads to a dramatic increase of pre-EEs and EEs, but maintains a population of ISCs (A). The ISC fraction progressively increases by co-expression of *da* (B) and tethered *da:da* (C), at the cost of EE differentiation (B, C) and pre-EE formation (C). **D.** Cell composition of GFP^+^ tissue from A-C. **E-F.** Knock- down of *Notch* with *esg^TS^-FO* results in excess of EEs and ISCs (E); simultaneous overexpression of *da:da* rescues both phenotypes (F). Solid and empty arrowheads: pre-EEs and ISCs, respectively; asterisks: EEs. Panel E is reproduced in **Figure S7A** to aid comparison. DAI: days after induction. Scale bars: 20µm. *p*-values (binomial regression): ● < 0.05; ●● < 0.01; ●●● < 0.001. See **Tables S1-2** for statistical details.

Loss of Notch leads to formation of masses of Pros-positive, EE-like cells (Bardin et al., 2010; Ohlstein and Spradling, 2006; **Figure 2E**). We further tested whether Da homodimers antagonise EE differentiation by co-expressing *da:da* and *Notch^RNAi^* with *esg^TS^-FO*, and found that Pros-positive appeared isolated and in low numbers (**Figure 2F**). Thus, Da:Da and Da:Sc oppose each other in EE differentiation; intriguingly, they seem to collaborate to induce proliferation (**Figure S2D-G**) and the formation of Dl- positive, Pros-negative ISCs (**Figure 2A-D**).

### Da:Da and Da:Sc collaborate to promote ISC identity

To understand how Da:Da and Da:Sc activate antagonistic transcriptional programs, we performed mRNA-seq analysis of purified *esg-Gal4 UAS-GFP* ISCs and EBs that overexpressed either *da^RNAi^*, *da*, *da:da* or *sc*. All conditions gave distinct transcriptional signatures (**Figures 3A** and **S3A** and **Table S3**). Interestingly, ∼1/3 of the genes downregulated upon *da^RNAi^* expression were common to those on *da* overexpression (**Figure 3A**). To explore these transcriptional signatures further, we looked at 57 cell type marker genes for ISCs, EBs, pECs and EEs (**Table S4**), and found that both loss and gain of Da downregulate EE-specific genes (**Figures 3B** and **S3C**). This may reflect that Da homodimers prevent EE differentiation (**Figures 1G-I** and **2B-D**) while Da:Sc dimers induce it (Li et al., 2017; **Figure S2A-C**). Consistent with this, overexpression of *sc* strongly induced EE-specific genes; it also increased expression of the ISC-specific genes *spdo* and *Dl*. Overexpression of *da:da* or *da* alone induced ISC-specific genes (*mira*, *spdo*), while most genes expressed in the absorptive lineage (*myo31DF*, *nub/pdm1*, *αTry*, *βTry*; *E(spl)mβ-HLH*, *E(spl)m3-HLH*, and other *E(spl)* genes outside the 57-gene panel) were upregulated in *da^RNAi^* overexpression and downregulated in the other conditions (**Figures 3B** and **S3B-C**). Thus, the transcriptome analysis supports our histofluorescence observations.

**Figure 3.**
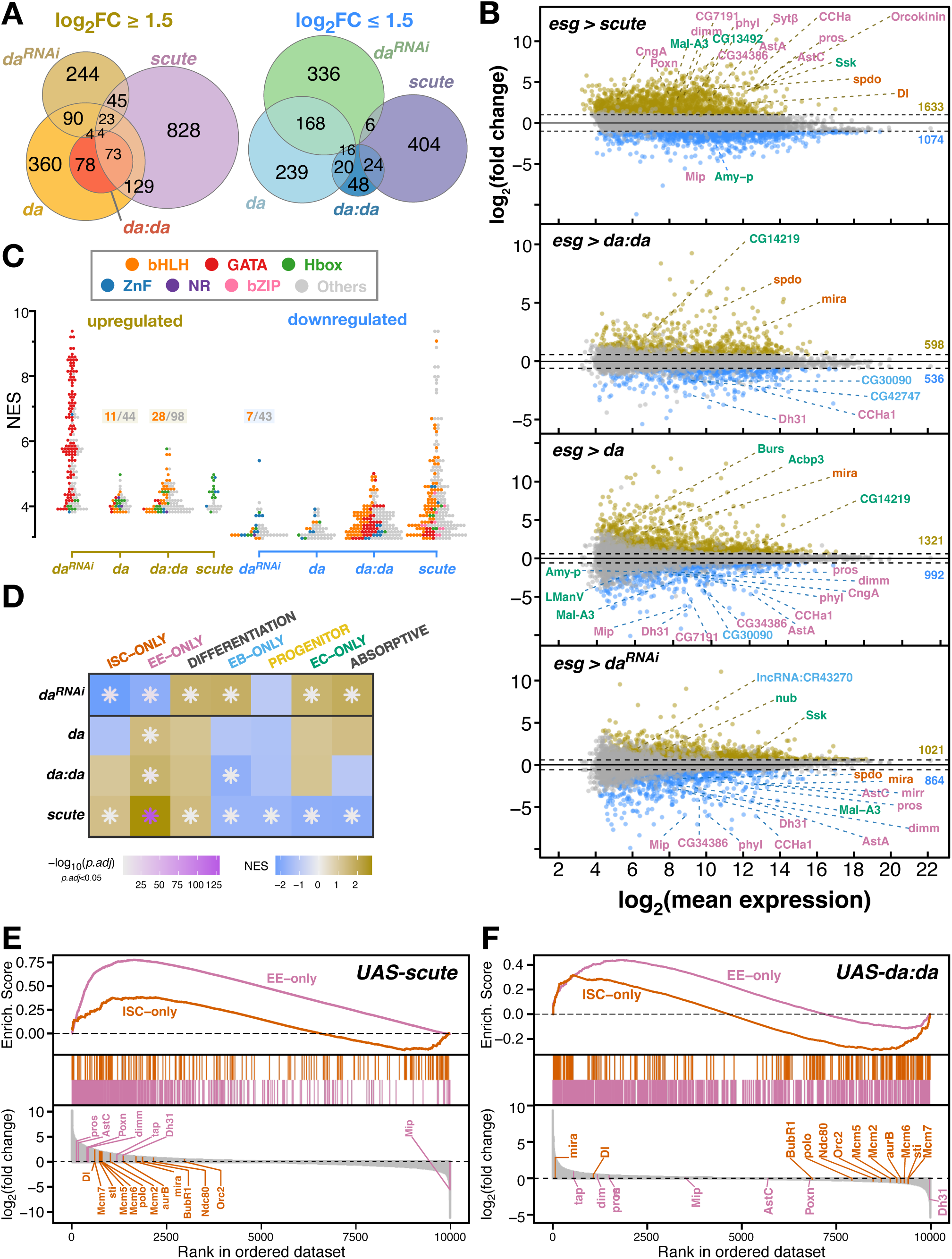
Da and Sc cooperate to induce the ISC transcriptional signature. **A.** Euler diagram with the sizes of gene sets differentially expressed (at│log_2_(FC)│≥1.5), and their approximate intersections, upon *da^RNAi^*, *da:da*, *da* and *sc* overexpression. **B.** MA plots for *esg^TS^ > sc*, *da:da*, *da* and *da^RNAi^*, respectively. Cell type markers are shown in colours matching Figure 1I. **C**. Normalised enrichment scores (NES) of DNA motifs found in differentially expressed genes (│log_2_(FC)│≥1.5) in the four conditions analysed. Dots represent individual motifs, coloured by the transcription factor family that binds them. Some swarms show their bHLH motif fraction. **D**. NES heatmap for cell-type specific gene sets; coloured asterisks indicate significance. **E-F.** Enrichment plots of the transcriptional profiles induced by *sc* (E) or *da:da* (F) for ISC- and EE-specific genes. *sc* induces a clear EE signature. Some ISC genes involved in replication (*Orc2*, *Mcm2*, *5*, *6* and *7*) and mitosis (*BubR1*, *polo*, *aurB*, *sti*, *Ndc80*) are repressed by *da:da* but activated by *sc*.

To determine whether the regulated genes were potential direct targets, we scored predicted regulatory elements close to differentially expressed genes for transcription factor binding motifs. We found E-boxes and other bHLH binding sites over-represented in upregulated genes upon *da* or *da:da* overexpression and downregulated ones upon *da^RNAi^* overexpression (**Figure 3C** and **Table S5**).

Gene Set Enrichment Analysis (GSEA) showed that *da^RNAi^* induced genes expressed in the absorptive lineage while reducing the expression of ISC- and EE-specific genes.

Meanwhile, *sc* overexpression did the opposite, leading to stronger expression of ISC-specific genes than *da:da* overexpression (**Figure 3D**). We looked at individual genes within the regulated, ISC-specific genes. While overexpression of either *da:da* or *sc* induced regulatory genes such as *Dl* or *mira*, they had opposite effects on genes encoding factors involved in DNA replication (*Orc2*, *Mcm2*, *Mcm5*, *Mcm6* and *Mcm7*) or mitosis (*polo*, *aurB*, *BubR1* and *Ncd80*): these were repressed by *da:da* but induced by *sc* (**Figure 3E-F**). This highlights the capacity of *sc* to regulate key ISC-specific genes.

We also found differences in broad functional annotations between the overexpression of *da^RNAi^* and that of *da*, *da:da* or *sc*. While loss of Da induces genes involved in metabolism, biosynthesis and energy storage and consumption, Da, Da:Da and Sc reduced the expression signatures of these processes and favoured signalling and regulatory genes (**Figure S3D-E**). We conclude that Da:Da and Da:Sc induce distinct signatures that promote the ISC and EE identities, respectively, and repress the active metabolism typical of EC function. However, Da:Da represses ISC-specific genes involved in replication and mitosis, which are upregulated by Da:Sc.

### Sc and Da can impart ISC properties

The transcriptional effects of *da:da* and *sc* overexpression on ISC-specific genes prompted us to evaluate the capacity of *sc*, *da* or *da:da* to impose ISC properties on more differentiated cells. We targeted EB, which are lineage-committed and postmitotic, using the driver *NRE^TS^-FO* (**Figure S4A**). Wild-type EBs labelled with *NRE^TS^-FO* either remained undifferentiated or became ECs; negligible numbers expressed Dl or Pros (**Figure 4A, E**). Driving expression of *sc*, *da* or *da:da* with *NRE^TS^-FO* abolished EC differentiation and led to a significant increase of Dl-positive cells (**Figure 4B-E**). *sc* overexpression in EBs also induced many pre-EEs and EE cells (**Figure 4D-E**). This suggested that Da, Da:Da and Sc could force EBs to revert to the ISC fate and, in the case of Sc, also to the secretory lineage. We evaluated whether these Dl-positive cells could, like ISCs do, undergo mitosis. We found mitotic (PH3-positive) cells within the EB-derived Dl-positive cells overexpressing *sc*; their proportion was of the same order as the mitotic, endogenous ISCs within the same tissue (2% vs 7%; **Figure 4F-G**). To assess whether *sc* may be promoting ISC properties in normal homeostasis, we analysed published single-cell RNAseq (scRNAseq) datasets (Hung et al., 2020; Li et al., 2022) (**Figure S4B**), and found that cells classified as ISC/EBs that expressed *sc* had on average much higher levels of *Dl* (**Figure 4H**), suggesting that *sc*, apart from inducing EE differentiation, normally promotes expression of ISC features.

**Figure 4.**
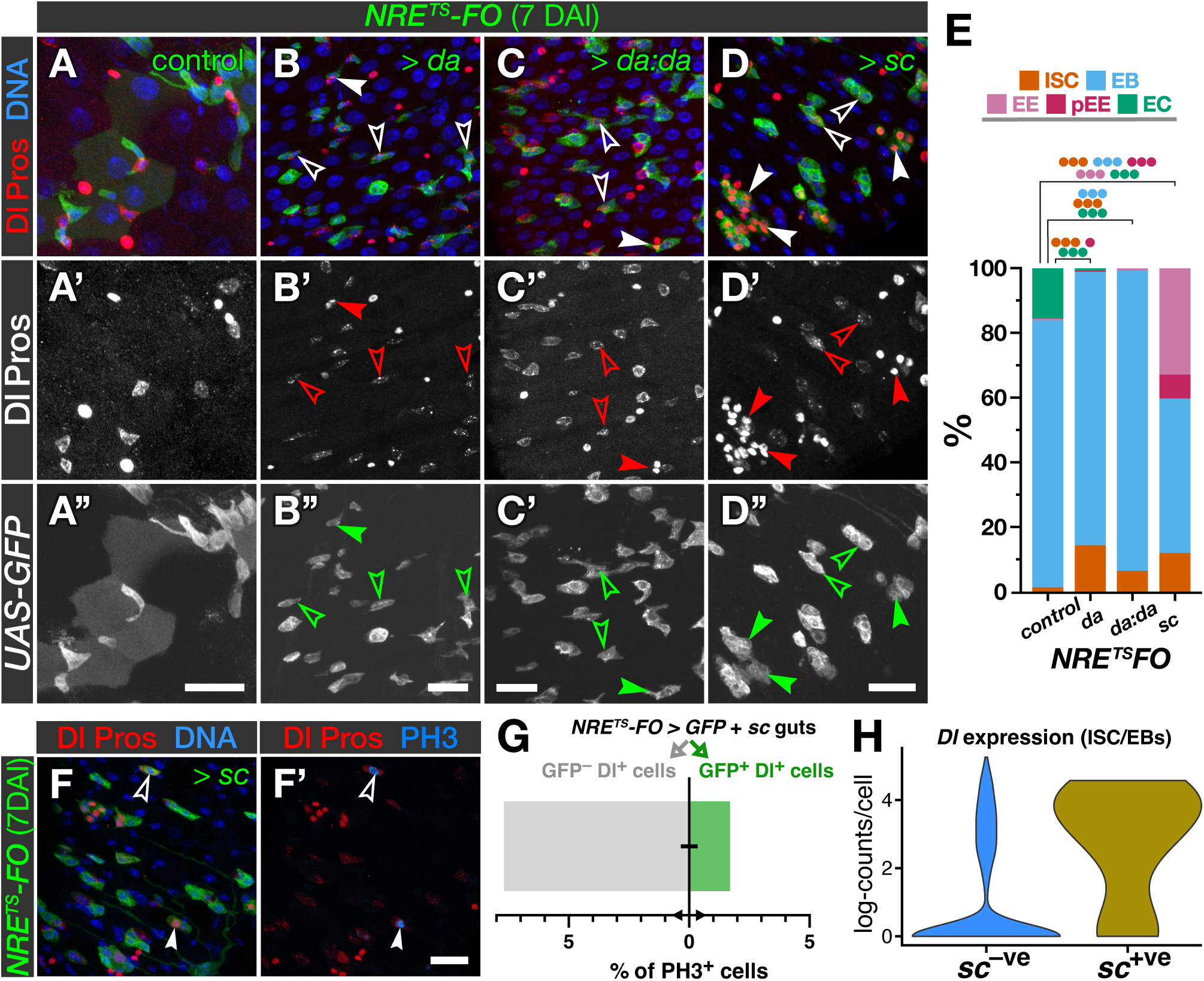
Da:Sc and Da:Da can impose ISC properties on EBs. **A-D.** Expression of either *da* (B), *da:da* (C) or *sc* (D) with *NRE^TS^-FO* (see **Figure S4A**) blocks normal EC differentiation (compare with A) and promotes re-expression of Dl (empty arrowheads) and EE differentiation (solid arrowheads). EB-derived EEs occur occasionally upon *da* or *da:da* overexpression but are very frequent upon *sc* overexpression, which also induces the formation of pre-EEs. **E.** Cell composition of GFP^+^ tissue from A-D. **F.** Overexpression of *sc* with *NRE^TS^-FO* shows both Dl^+^ and Pros^+^ cells (empty and solid arrowheads, respectively) undergoing mitosis (phospho- H3^+^). **G.** Proportions of mitotic Dl^+^ cells (Pros^+^ or Pros^—^) within and out of the population of cells co-expressing *GFP* and *sc* in *NRE^TS^-FO>GFP* + *sc* intestines (N=303 cells). **H.** Expression of *Dl* found in ISCs/EBs by scRNAseq (see **Figure S4B**), segregated by the expression of *scute* (zero vs non-zero counts). DAI: days after induction. Scale bars: 20µm. *p*-values (binomial regression): ● < 0.05; ●● < 0.01; ●●● < 0.001. See **Tables S1-2** for statistical details.

### Emc promotes EC differentiation by titrating Da

Da and Sc functions are often antagonised through direct binding and titration by the HLH class V factor Emc, which prompted us to test *emc* function in intestinal homeostasis. Using the protein trap line *emc^CPTI002740^*(Lowe et al., 2014), we found *emc* expressed in all cell types of the adult gut but predominantly by EBs and ECs (**Figures 5A-C**). scRNAseq data (Hung et al., 2020; Li et al., 2022) show that *emc* expression is mainly in EBs and ECs of the posterior midgut, with cell type distributions similar to bona-fide markers of posterior midgut ECs (pECs) and EBs (**Figure S5A-H**). Consistent with this, *emc* expression decreases along the transcriptional trajectory from ISC/EB to EE and increases in the ISC/EB to pEC trajectory (**Figures 5D** and **S5I**). MARCM clones mutant for *emc^LL02590^* (**Figure S5J**) had fewer differentiated cells and were enriched in Dl-positive ISCs (**Figure 5E-F**, **J**). Clones for alleles *emc^1^* and *emc^AP6^* (**Figure S5J**) showed similar enrichment in Dl-positive ISCs compared to wild-type clones (**Figure 5G-J**). Hypomorphic viable *emc^LL02590^/emc^1^* and *emc^LL02590^/emc^P5C^*heterozygotes (**Figure S5J**) showed an increase in ISCs/EBs at the expense of ECs, accompanied by higher levels of Dl in ISCs and its ectopic expression in ECs, compared to control *emc^LL02590^/+* guts (**Figure 5K-N**). Knockdown of *emc* with *esg^TS^-FO* phenocopied the overexpression of *da*: ISCs/EBs gave rise to labelled clusters of similar size to the controls (**Figure 1L**) where differentiation was severely impaired (**Figure 5O-Q, S**). The strength of the effect on differentiation in the *emc^RNAi^* transgenes correlated with their efficacy in depleting Emc protein and inducing *extra macrochaetae* phenotypes (**Figure S5K-R**). The strongest *emc^RNAi^* (*NIG-1007R*) induced, as with *da* overexpression, the loss of many *esg^+^* cell nests by apoptosis (as death was prevented by co-expression of p35; **Figure S6A-B**). We conclude that *emc* is necessary for EC differentiation and the survival of ISCs and EBs. Overexpression of *emc* with *esg^TS^-FO* resulted in all ISCs/EBs differentiating into ECs (**Figure 5R-S**), as expected (Lan et al., 2018). Therefore, Emc is both necessary and sufficient to direct absorptive differentiation.

**Figure 5.**
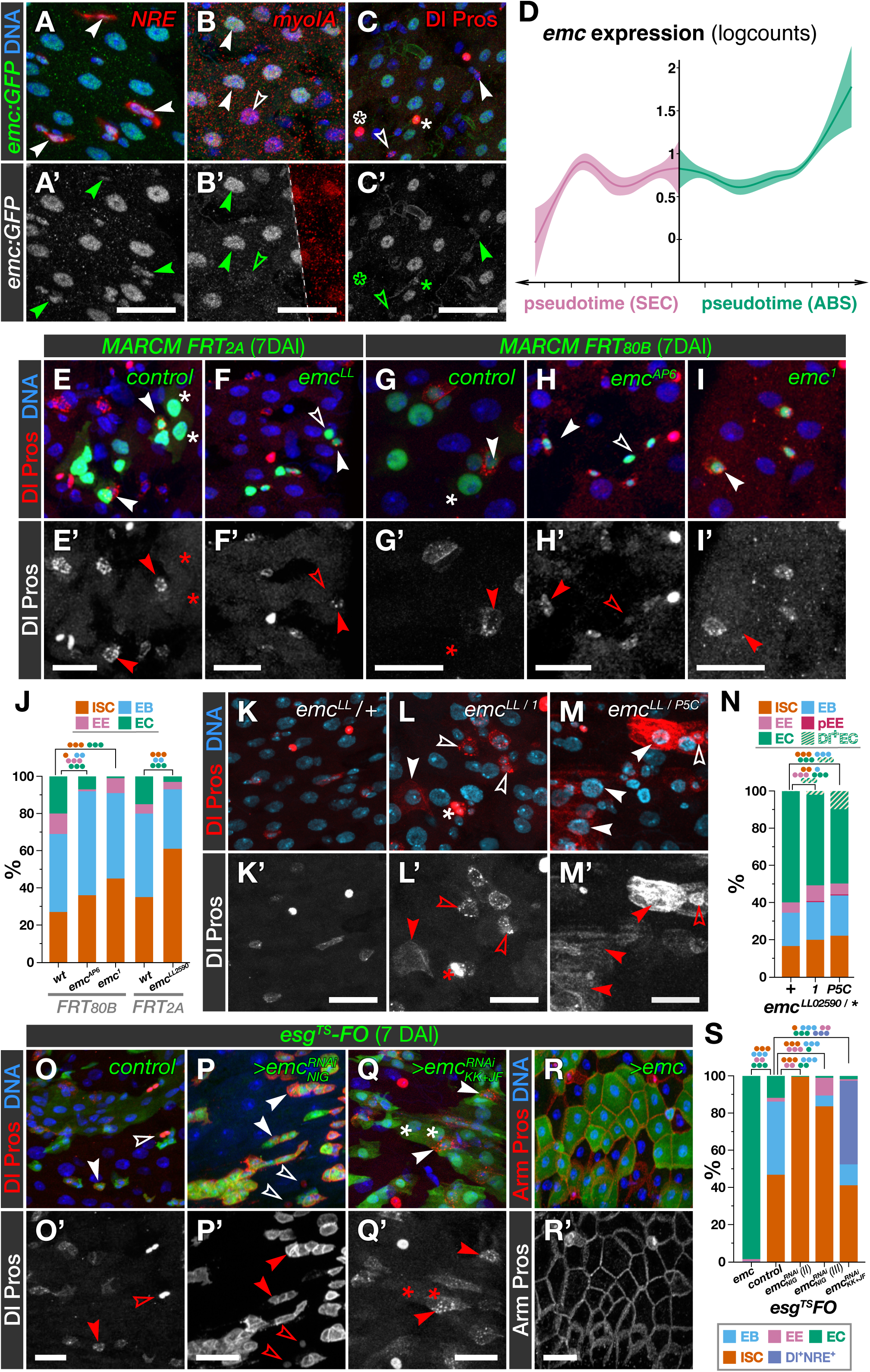
*emc* is necessary and sufficient to induce EC differentiation. **A-C.** Expression of Emc-GFP in the homozygous viable protein-trap insertion *emc^CPTI002740^*. Emc is expressed in EBs (*NRE-lacZ*^+^, arrowheads) (A) and some ECs (*myoIA-lacZ*^+^, solid arrowheads) but not all (empty arrowheads) (B). Some ISCs (Dl^+^, solid arrowheads) and EEs (Pros^+^, solid asterisks) express Emc, but many do not (empty arrowheads and asterisks) (C). **D.** *emc* expression along the pseudotime trajectories from ISC/EB cells into EEs and ECs (**Figure S5I**) decreases towards the secretory fate and increases towards the absorptive one. **E-I.** Cells in MARCM clones for *emc^LL02590^* (F), *emc^AP6^* (H) and *emc^1^* (I) are enriched in ISCs (solid arrowheads) and EBs (empty arrowheads) compared to controls (E and G). *FRT_80B_* and *FRT_2A_* are two independent insertions for inducing MARCM clones in chromosomal arm 3L. **J.** Cell composition of clones from E-I. **K-M.** Midguts of viable hypomorphic mutants *emc^LL02590/1^* (L) and *emc^LL02590/P5C^* (M) contain more ISCs, preEEs (L, asterisk) and EEs, and less ECs, than heterozygous controls (K). In these mutants, Dl expression is elevated in some ISCs (L, M, empty arrowheads) and ectopic in some ECs (L, M, solid arrowheads). **N.** Cell composition in genotypes from K-M. **O-Q.** Expressing *emc^RNAi^* with *esg^TS^-FO* using transgenes *NIG-1007R* (P) or *KK108316* and *JF02300* together (Q) increases the proportion of Dl^+^ cells and EEs (empty arrowheads) and reduces that of ECs (solid arrowheads) compared to controls (O). *emc^RNAi^* also increases Dl expression (P, Q), especially with *NIG-1007R* (P). **R.** Overexpression of *emc* with *esg^TS^-FO* forces differentiation into ECs. **S.** Cell composition in genotypes from O-R. DAI: days after induction. Scale bars: 20µm. *p*-values (binomial regression): ● < 0.05; ●● < 0.01; ●●● < 0.001. See **Tables S1-2** for statistical details. Panels F, P, R are reproduced in Figures 6D**, S7J, G**, respectively, to aid comparison.

If Emc promoted EC differentiation by titrating Da and Sc, *emc* function should depend on *da*. We tested this by generating MARCM clones mutant for *da* and expressing *emc^RNAi^* or mutant for *emc* and expressing *da^RNAi^*. Both conditions led to EC differentiation like in *da^10^* clones (**Figure 6A-C**, compare with **6D**). Considering our previous results where *sc* promoted ISC characteristics (**Figures 3** and **4**) and that ISCs express Sc (Chen et al., 2018; Doupé et al., 2018), we tested whether the effects of *emc* loss depended on *sc* or any other member of the *AS-C*. We induced MARCM clones for *Df(1)sc^B57^* (a deletion of the *AS-C*; González et al., 1989) expressing *emc^RNAi^* and compared them with clones that only expressed *emc^RNAi^*. Differentiation was impaired in both conditions (**Figure 6F-G**), but Dl expression was four-fold lower in the absence of the AS-C (**Figure 6H**). We conclude that Emc promotes EC differentiation and dampens Dl expression by preventing formation of Da:Da and Da:Sc dimers, respectively.

**Figure 6.**
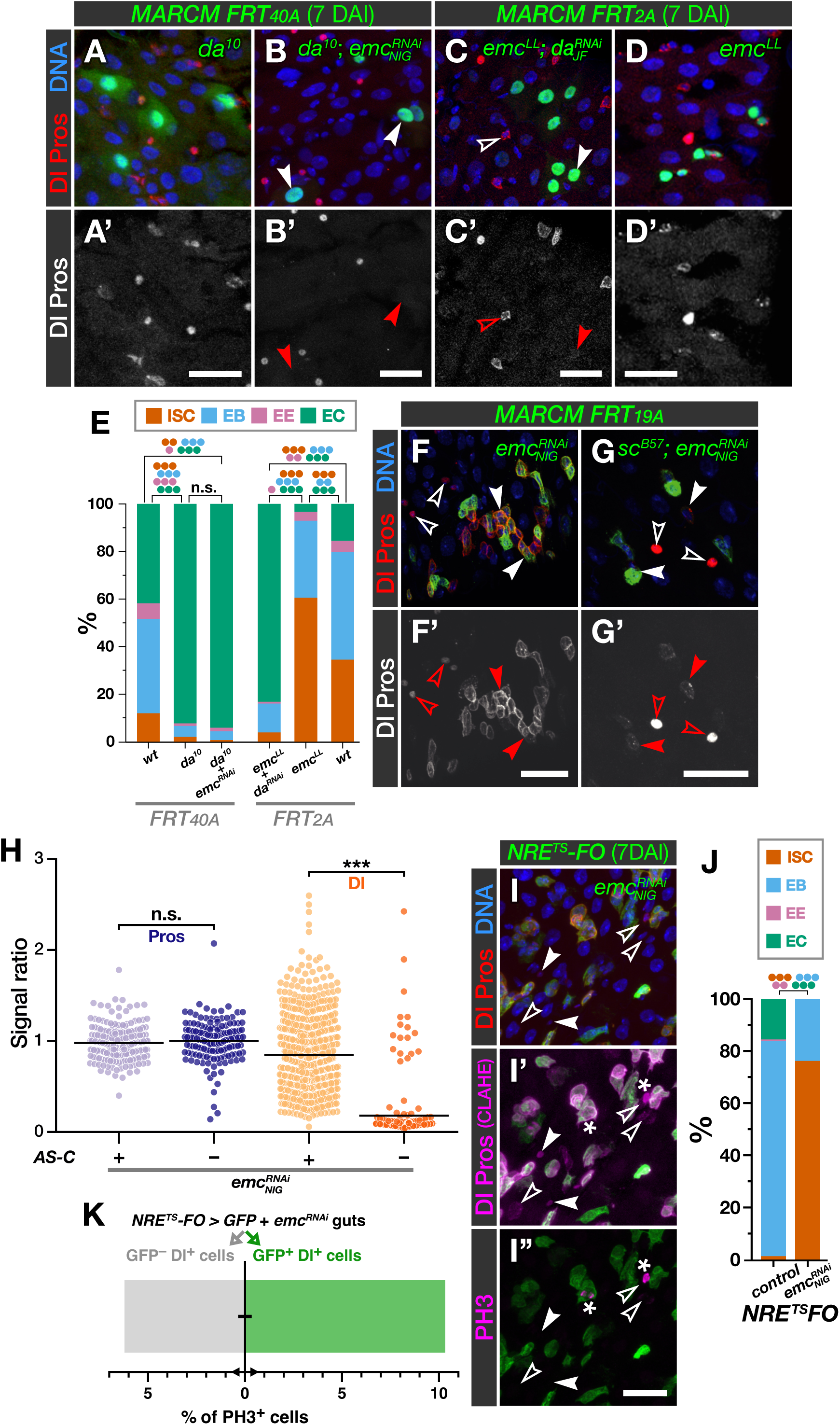
*emc* antagonises *da* and *sc* in the intestine and maintains EBs committed. **A-D.** *da* is epistatic over *emc*. Solid arrowheads: ECs; empty arrowheads: ISCs. Cells in MARCM clones that are mutant for *da^10^*and express *emc^RNAi^ NIG-1007R* (B) mostly differentiate into ECs, like in *da^10^* clones (A). Cells in MARCM clones that are mutant for *emc^LL02590^* and express *da^RNAi^ JF02488* (C) also differentiate into ECs, suppressing the reduced differentiation of *emc^LL02590^* cells (D). **E.** Cell composition of clones from A- D and Figure 1I. Note that *da^10^* is statistically indistinguishable from *da^10^*; *emc^RNAi^*. **F-G.** Emc antagonises Sc in inducing *Dl*. Loss of the *achaete/scute-Complex* (*AS-C*) using deficiency *Df(1)sc^B57^* reduces the elevated Dl expression observed with *emc^RNAi^* (G, compare with F). **H.** Quantification of relative levels of Dl per cell, normalised by Pros (see Methods), for 210, 146, 718 and 210 cells per group from left to right. Horizontal lines are averages per category. *p*-values (Mann-Whitney test): ⁕⁕⁕ < 0.001; n.s., ≥ 0.05. **I-K.** Expressing *emc^RNAi^ NIG-1007R* with *NRE^TS^-FO* prevents EB differentiation and activates Dl expression (I’, solid arrowheads) at higher levels than extant ISCs (arrowheads; extant ISCs are detectable after contrast-limited adaptive histogram equalization, CLAHE). These cells are mitotic (I’’, phospho-Histone H3, asterisks) at levels similar to extant ISCs (K) and have presumably reverted into ISCs (quantified in J). Scale bars: 20µm. *p*-values (binomial regression in stacked bars): ● < 0.05; ●● < 0.01; ●●● < 0.001; n.s. ≥ 0.05. See **Tables S1-2** for statistical details. Panels A, D are reproduced from Figures 1B**, 5F**, respectively, to aid comparison.

### *emc* is required for the commitment of EBs

The requirement of *emc* for differentiation prompted us to investigate whether it was necessary to maintain the commitment of EBs as absorptive progenitors. We used the EB-specific driver *NRE^TS^-Gal4*, which never co-expresses with Dl in the wild-type, to express *emc^RNAi^*. This led to Dl expression in *NRE^+^* cells (**Figure S6C-D**), suggesting that *emc*-depleted EBs may revert to ISCs. We verified this with *NRE^TS^-FO* driving *emc^RNAi^* expression and observed that *emc*-depleted EBs give raise to Delta-positive cells (**Figure 6I-J**) that undergo mitosis (PH3-positive) at similar frequency to neighbouring normal ISCs (**Figure 6I, K**). This suggests that *emc* is necessary for the commitment of EBs.

### Da/Emc regulate ISC date independently of Notch

Notch signalling is key for ISC differentiation and absorptive fate acquisition (Micchelli and Perrimon, 2006; Ohlstein and Spradling, 2006). We found that simultaneous knock-down of *N* and *da* with *esg^TS^-FO* prevented the tumorous expansion of Dl-positive and Pros-positive cells typically found with *N^RNAi^* alone (Micchelli and Perrimon, 2006; Patel et al., 2015) and led to ECs differentiation (**Figure S7A-B**). This suggests that Notch could be regulating *da*. To activate or prevent Notch signalling, we used *esg^TS^-FO* to knock-down or overexpress Hairless (H), the specific co-repressor for the transcriptional targets of Notch (reviewed in Bray and Furriols, 2001).

Overexpression of *H* with *esg^TS^-FO* resulted in expansion of Dl-positive cells and EE differentiation (**Figure 7A, I**; Bardin et al., 2010). Simultaneously knocking down *da* significantly induces EC differentiation, but not to the levels induced by *da* knock-down alone (**Figure 7B, I**). Conversely, expression of *H^RNAi^* with *esg^TS^-FO* leads to an increase in EC differentiation (**Figure 7C**; Bardin et al., 2010) and the expression of *NRE-lacZ* in all diploid cells (**Figure 7I**). The co-expression of *da:da* with *H^RNAi^* prevents all EC differentiation, but extends *NRE-lacZ* expression to all *esg^+^* cells, at the expense of Dl (**Figure 7D, I**). Thus, simultaneously inducing or interfering with *da* and Notch signalling leads to phenotypes in between their respective individual manipulations. This suggests that *da* and Notch signalling can act in parallel.

**Figure 7.**
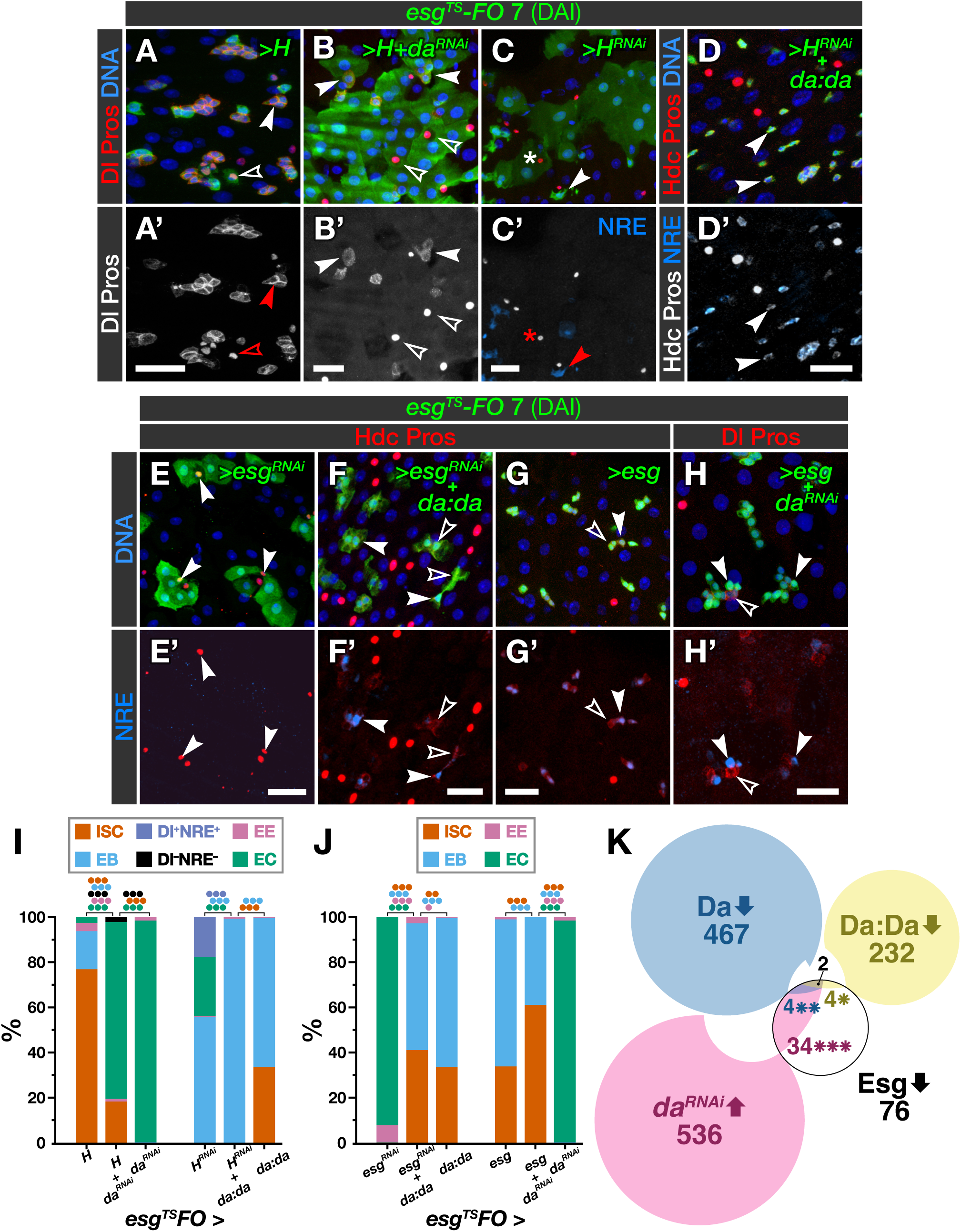
*da* is epistatic to Notch signalling and acts in parallel with *esg*. **A-B.** Expressing *H* with *esg^TS^-FO* (A) blocks EC differentiation and leads to accumulation of ISCs. This is rescued by simultaneous expression of *da^RNAi^* (B). Solid arrowheads: ISCs; empty arowheads: EEs. **C-D**. Expressing *H^RNAi^* with *esg^TS^-FO* (C) induces EC differentiation, some EE differentiation (asterisk) and the expression of the *NRE-lacZ* reporter in all *esg^+^* cells (solid arrowhead). Simultaneous expression of *da:da* (D) blocks the formation of ECs, as in *esg^TS^-FO>da:da* alone, and all cells become *NRE-lacZ^+^* (solid arrowheads). **E-F.** Knock-down of *esg* with *esg^TS^-FO* (E) induces differentiation of all cells into ECs and EEs (solid arrowheads); this is prevented by simultaneous expression of *da:da* (F), which blocks differentiation of ISCs (empty arrowheads) and EBs (solid arrowheads) to the same levels as in *esg^TS^-FO>da:da* alone. **G-H.** Expression of *esg* with *esg^TS^-FO* (G) blocks differentiation regardless of whether *da* is simultaneously knocked-down (H). Empty arrowheads: ISCs; solid arrowheads: EBs. DAI: days after induction. Scale bars: 20µm. **I-J.** Cell composition of GFP^+^ tissue from A-D and E-H, respectively; data from Figure 1I is replicated here to aid comparison. *p*-values (binomial regression for individual cell types): ● < 0.05; ●● < 0.01; ●●● < 0.001. See **Tables S1-2** for statistical details. **K.** Modified Euler diagram showing the overlap between gene sets repressed directly by Esg (white) with those either repressed by overexpression of *da* (blue) or *da:da* (yellow) or de-repressed by expression of *da^RNAi^*(red), all three at│log_2_(FC)│≥1). Areas approximate set sizes and their intersections. Overlaps between *da-*regulated sets are not depicted. *p*-values for overlap sizes (Esg∩Da, Esg∩Da:Da and Esg∩*da^RNAi^*; hypergeometric test with Bonferroni correction) are: ⁕⁕⁕ < 1e–36, ⁕ < 0.01, ⁕ < 0.05.

*emc* transcription is regulated by Notch signalling in other tissues (Adam and Montell, 2004; Baonza et al., 2000; Baonza and Freeman, 2001; Bhattacharya and Baker, 2009; Spratford and Kumar, 2015), and we found that co-expressing *emc* and *N^RNAi^* with *esg^TS^-FO* leads to EC differentiation (**Figure S7C**). Therefore, it is possible Notch could induce EC differentiation at least partially through *emc*. We found that *emc* expression induces EC differentiation rapidly (Puig-Barbe, 2018), so these *esg^TS^-FO*>*emc*+*N^RNAi^*ECs could have differentiated before N protein was depleted. To inhibit Notch signalling faster, we co-expressed *emc* and *H* with *esg^TS^-FO*. This led to an increase in EC differentiation compared to expression of *H* alone, but far from the near-complete EC differentiation observed when only expressing *emc* (**Figure S7D-G**). This suggests that Emc can induce differentiation in the absence of Notch signalling, but not efficiently. Moreover, expressing *H^RNAi^* and *emc^RNAi^*with *esg^TS^-FO* did not prevent the excess EC differentiation observed upon expression of *H^RNAi^* alone; this contrasts with the capacity of *emc^RNAi^* to block differentiation entirely (**Figure S7D, H-J**). Therefore, Emc is dispensable for Notch to induce EC differentiation. Moreover, clonal expression of the Notch intracellular domain does not induce *emc* expression in any cell type of the midgut epithelium (**Figure S7K**). We conclude that *emc* and Notch signalling act independently to promote the absorptive fate.

### Da:Da and Esg block differentiation independently

Esg, a transcription factor of the Snail family, regulates ISCs by preventing ISC/EB differentiation (Korzelius et al., 2014; Loza-Coll et al., 2014). As Da:Da dimers have the same capacity, we considered whether Esg and Da act together to maintain stemness. Expression of *esg^RNAi^* with *esg^TS^-FO* led to differentiation into ECs and EEs (**Figure 7E, J**), as expected (Korzelius et al., 2014; Li et al., 2017; Loza-Coll et al., 2014).

Simultaneous overexpression of *esg^RNAi^* and *da:da* blocked EC differentiation and significantly reduced formation of EEs (**Figure 7F, J**). In turn, expression of *esg* with *esg^TS^-FO* blocked differentiation, irrespective of whether *da^RNAi^* was also expressed (**Figure 7G-H, J**). This independence seems mirrored in their down-regulated gene sets: almost half the Esg repression targets overlap with genes downregulated by misexpression of *da* or *da:da* or upregulated by misexpression of *da^RNAi^*, including genes essential for EC function like *nub/pdm1*, *ssk* and *Tsp2A* (**Figure 7K**). Finally, we observed that expression of neither *da* nor *esg* were affected by the overexpression the other (**Figure S7L-M**). We conclude that *da* and *esg* contribute to stemness independently of each other.

## DISCUSSION

We demonstrate a central role for a bHLH factor code in the acquisition and maintenance of three alternative cell fates in the adult *Drosophila* intestine. Class I homodimers (Da:Da) maintain the progenitor state of ISCs/EB, with changes in dimerization partners governing the fate transitions. Sequestration of Da with class V HLH factor Emc into Da:Emc dimers incapable of DNA binding induces progenitor cells to acquire the absorptive fate, while formation of Da:Sc dimers by binding of Da with class II bHLH Sc initiates EE differentiation (**Figures 2** and **S2**; Bardin et al., 2010; Zeng and Hou, 2015). Moreover, Emc is required in EBs to maintain their committed state, while low levels of Sc boost aspects of the ISC transcriptional state.

### Three cell fates regulated by a dimerization network

Networks involving class I, II and V bHLH factors regulate the development of the *Drosophila* retina (Bhattacharya and Baker, 2011) and peripheral nervous system (Cubas et al., 1991; Troost et al., 2015; Van Doren et al., 1991). However, in these cases, the choice is between only two alternative fates (neural vs epidermal), with Da homodimers promoting the same fate as heterodimers between Da and a class II bHLH proneural factor (Bhattacharya and Baker, 2011; Li and Baker, 2018; Troost et al., 2015). By contrast, in the adult midgut Da:Da and Da:Sc promote distinct fates (progenitor and secretory, respectively) through distinct transcriptional programs while Emc titrates both dimers to allow EC differentiation. This integrated mechanism suggests that the balance between ISC self-renewal and absorptive or secretory differentiation rests on a triple choice, rather than two consecutive binary decisions. An equivalent network could operate in similar stem cell systems, and it is tempting to speculate of this possibility in the mammalian intestine, where the relevant factors (mouse *da* homologs *E2a* and *Heb*, *emc* homolog *Id1* and *sc* homolog *Acsl1*) are expressed in the crypts of Lieberkühn and have roles in fate determination (van der Flier et al., 2009; Wice, 1998; Zhang et al., 2014).

### A balance of factors regulates intestinal stem cell fate

We refer to the Da/Emc/Sc network as a bHLH ‘code’ due to its modularity, but it does not behave as a Boolean switch; the relative abundance of the different components seems critical. Da, the centrepiece of the network, forms homodimers that maintain stemness and prevent EC and EE differentiation (**Figures 1** and **2**), but it is also an essential partner for Sc in EE differentiation (**Figure S2**). The activity of Da homodimers in promoting stemness is also nuanced, as in parallel to activating some ISC genes and repressing some EE/EC ones, Da:Da downregulates proliferation effector proteins, in agreement with its previously reported anti-proliferative activity (Andrade-Zapata and Baonza, 2014). Strikingly, an excess of Da monomers (whether induced by direct overexpression or indirectly by loss of Emc) promotes apoptosis (**Figures 1** and **S6**), while the excess of Sc or tethered Da:Da does not. It follows that Da also participates in an apoptosis-promoting complex yet to be identified. By contrast, the activity of Emc is determinant, but not intrinsically instructive: through titration of Da and Sc cells have to differentiate but lack the capacity to initiate EE differentiation, so they differentiate into ECs by default. However, Emc loss in EBs induces expression of ISC features, possibly through the increased activity of Sc (**Figure 6**), and prevents Da from inducing apoptosis of ISC/EBs (**Figure S6**). This makes Emc also an essential factor to maintain the progenitor population. Sc, which is normally expressed at low levels in ISCs (Chen et al., 2018; Doupé et al., 2018), triggers EE differentiation when expressed over a threshold (Amcheslavsky et al., 2014; Bardin et al., 2010; Chen et al., 2018). We find that Sc boosts expression of essential ISC genes, such as Dl or those involved in proliferation. Moreover, Sc seems to induce fate reversal in EBs. Therefore, Sc also boosts the ISC function, which sits well with the role of its homolog *Acsl2* in maintaining mammalian ISCs (van der Flier et al., 2009).

We conclude that a dynamic balance of Da, Emc and Sc is required for homeostasis, and this bHLH code is not composed not of ‘master’ but contextual regulators. This pushes the question towards the regulation of their protein concentrations and/or functions across intestinal cell types and over time.

In line with this view, other essential regulators of ISC fate operate in parallel to the bHLH network. Excess of either Esg or Da can compensate for the absence of each other to maintain ISCs/EBs undifferentiated, suggesting that at physiological levels, when either is essential for stemness (**Figure 1**; Korzelius et al., 2014), they have some additive effect. This may be reflected in the significant overlap of their downregulated genes (**Figure 7**), which includes the EC-fate inducer *nub/pdm1*. On the other hand, Esg seems to have a more specific role than Da:Da in preventing EE differentiation and lacks its anti-proliferative activity (Korzelius et al., 2014; Loza-Coll et al., 2014), suggesting that they also govern non-shared aspects of ISC identity and function. Similarly, both Notch signalling and Emc can induce EC differentiation in the absence of the other, which suggests that the absorptive fate does not simply arise by default when Da and Sc are not functioning. These multiple, non-redundant layers of regulation likely allow fine-tuning the ISC function.

### Multiple factors maintain enteroblast commitment

Notch signalling induces the formation of EBs (Micchelli and Perrimon, 2006; Ohlstein and Spradling, 2006), which under normal conditions give rise to mature ECs without further division (Wang et al., 2015; Yin and Xi, 2018; Zeng and Hou, 2015; **Figure 4**). EBs are relatively long-lived (Antonello et al., 2015), and though they are recognised by expression of the *NRE*, once formed they do not need Notch to maintain their commitment (Siudeja et al., 2015). However, we found that they require Emc, as its depletion in EBs gave rise to Dl-expressing proliferative cells–likely a reversal of fate towards ISC. Qualitatively similar effects resulted from mis-expressing *da*, *da:da* or *sc* in EBs. We interpret that combined baseline levels of Da and Sc in EBs may lead them to revert their fate into ISCs if not titrated by Emc. EB-to-ISC reversion is also prevented by transcription factor Sox21a (Zhai et al., 2015) and the global co-repressor Groucho (Guo et al., 2019). EB-to-EE trans-differentiation has also been observed, when Ttk69 or Klu were depleted, or Phyllopod overexpressed, in EBs (Korzelius et al., 2019; Wang et al., 2015; Yin and Xi, 2018). These observations underscore the plasticity of the EB and resemble the behaviour of EC precursors in the mammalian intestine, which can dedifferentiate and repopulate the intestinal crypt during regeneration (Tetteh et al., 2016).

### Limitations of the study

Our interpretation of the functional relationships between Da, Sc and Emc are based on misexpression tools with no control over resulting stoichiometry. Future studies should address this limitation with newly developed tools with more precise control (Goldner et al., 2023; Yesbolatova et al., 2020; Diao and White, 2012). Our analysis of *emc* function involved the *emc^RNAi^* transgene *NIG-1007R*, which is far more effective at depleting Emc than either *KK108316* or *JF02300* or both combined (**Figure S5**).

Several *emc* loss of function conditions affecting large amounts of tissue induce increased Dl expression (**Figure 5**), but with *NIG-1007R* this effect is much higher and not observed in MARCM *emc* null clones. This may point to non-cell-autonomous, suppressing effects on Dl expression that are only inactivated when the whole tissue or the whole progenitor population are affected. We are confident that *NIG-1007R* elicits genuine *emc* loss phenotypes, because the strength of the increase in Dl levels correlates with the strength of the *emc^RNAi^* transgenes (**Figures 5** and **S6**); *NIG-1007R* phenocopies the overexpression of *da*, including the induction of apoptosis (which *KK108316* and *JF02300* cannot recapitulate; **Figures 1**, **5** and **S5**); we can fully rescue the increase in Dl expression induced by *NIG-1007R* by simultaneous loss of *sc* (**Figure 6**). While our tenet does not hinge on this observation, it deserves future attention.

## ACKNOWLEDGEMENTS

We thank the anonymous reviewers for helpful suggestions, Nicholas Baker, Allison Bardin, Antonio Baonza, Sonsoles Campuzano, Sangbin Park, Mike Taylor, Shinya Yamamoto, Alfonso Martínez-Arias, the Bloomington *Drosophila* Stock Center, the Vienna *Drosophila* Resource Center, the *Drosophila* Genetics Resource Center (Kyoto) and the National Institute of Genetics (Japan) for providing fly stocks, the Developmental Studies Hybridoma Bank (University of Iowa) for supplying antibodies, and Juan Modolell, Sonsoles Campuzano, Catherine Hogan, Fernando dos Anjos-Afonso, Florian Siebzehnrubl, Terrence Trinca, Sonia López de Quinto, Helen White-Cooper, Mike Taylor and Wynand van der Goers van Naters for critical comments on the manuscript and useful discussions. We acknowledge the data analysis team of the College of Biomedical and Life Sciences of Cardiff University and the FACS and Imaging Core Facility at the Max Planck Institute for Biology of Ageing for technical assistance.

This work was supported by funding from Cardiff University to JdN and AP, from the University of Essex and NC3Rs SKT grant NC/W001047/1 to PVW and JdN, a DFG-Grant KO5594/1-1 and an EMBO Long-Term Fellowship to JK, a FAPESP fellowship #2021/00393-9 to VDN, a FAPESP São Paulo Excellence Chair #2019/16113-5 to PVW, and an ERC Advanced Grant no. 268515 to BAE.

JdN dedicates this work to the memory of Juan Modolell and Rosa María Aguilar.

## AUTHOR CONTRIBUTIONS

Conception of study, supervision: JdN. Data acquisition: AP, SD, SA, JK, HM and JdN. Analysis and interpretation: AP, VDN, JdN, JK. Training, supervision: JdN, JK, PVW, BE. Writing the manuscript: JdN, AP. Revising and approval of the manuscript: all authors.

