## Supplementary material for "A bHLH interaction code controls bipotential differentiation and self-renewal in the *Drosophila* gut": Methods

### STAR★Methods

#### KEY RESOURCES TABLE

| REAGENT or RESOURCE | SOURCE | IDENTIFIER |
|---|---|---|
| **Antibodies** | | |
| Mouse monoclonal anti-Delta, extracellular domain, c594.9b (1:50) | Developmental Studies Hybridoma Bank (DSHB) | RRID: AB_528194 |
| Mouse monoclonal anti-Prospero, MR1A (1:200) | DSBH | RRID: AB_528440 |
| Mouse monoclonal anti-Headcase, HDC U33 (1:100) | DSHB | RRID: AB_10659722 |
| Mouse monoclonal anti-Armadillo, N2 7A1 (1:100) | DSHB | RRID: AB_528089 |
| Rabbit polyclonal anti-Prospero (1:1000) | (Vaessin et al., 1991) | N/A |
| Guinea pig polyclonal anti-Emc (1:1000) | Antonio Baonza (CSIC, Spain) | N/A |
| Rabbit polyclonal anti-Phospho-Histone H3 (Ser10) (1:200) | Cell Signalling | Cat# 9701, RRID: AB_331535 |
| Rabbit polyclonal anti-GFP (1:2000) | Abcam | Cat# ab6556, RRID: AB_305564 |
| Chicken polyclonal anti-GFP (1:3000) | Abcam | Cat# ab13970, RRID: AB_300798 |
| Chicken polyclonal anti-beta Galactosidase (1:2000) | Abcam | Cat# ab9361, RRID: AB_307210 |
| Rabbit polyclonal anti-beta Galactosidase (1:10000) | ThermoFisher Scientific | Cat# A11132, RRID: AB_221539 |
| Goat anti-Mouse IgG, Alexa Fluor 568-conjugated (1:500) | ThermoFisher Scientific | Cat# A-11031, RRID: AB_144696 |
| Donkey anti-Mouse IgG, Alexa Fluor 594-conjugated (1:500) | ThermoFisher Scientific | Cat# A-21203, RRID: AB_2535789 |
| Goat anti-Mouse IgG, Alexa Fluor 633-conjugated (1:500) | ThermoFisher Scientific | Cat# A-21050, RRID: AB_2535718 |
| Goat anti-Rabbit IgG, Alexa Fluor 488-conjugated | ThermoFisher Scientific | Cat# A-11008, RRID: AB_143165 |
| Donkey anti-Rabbit IgG, Alexa Fluor 594-conjugated | ThermoFisher Scientific | Cat# A-21207, RRID: AB_141637 |
| Goat anti-Rabbit IgG, Alexa Fluor 633-conjugated | ThermoFisher Scientific | Cat# A-21070, RRID: AB_2535731 |
| Goat anti-Guinea pig IgG, Alexa Fluor 546-conjugated | ThermoFisher Scientific | Cat# A-11074, RRID: AB_2534118 |
| Goat anti-Chicken IgY, Alexa Fluor 488-conjugated | ThermoFisher Scientific | Cat# A-11039, RRID: AB_2534096 |
| Goat anti-Chicken IgY, Alexa Fluor594-conjugated | ThermoFisher Scientific | Cat# A-11042, RRID: AB_2534099 |
| Goat anti-Chicken IgY, Alexa Fluor 633-conjugated | ThermoFisher Scientific | Cat# A-21103, RRID: AB_2535756 |
| **Chemicals, peptides, and recombinant proteins** | | |
| Phosphate buffered saline tablets | Sigma-Aldrich | Cat# P4417 |
| Triton X-100 | Sigma-Aldrich | Cat# T8787 |
| Bovine Serum Albumin fraction V | Roche | Cat# 10735108001 |
| Formaldehyde solution | Sigma-Aldrich | Cat# F8775 |
| Hoechst 33342 (used at 2 µg/ml) | Sigma-Aldrich | Cat# B2261 |
| N-propyl-gallate | Sigma-Aldrich | Cat# 02370 |
| Glycerol (spectrophotometric grade) | Sigma-Aldrich | Cat# G9012 |
| **Critical commercial assays** | | |
| PureLink RNA mini preps | ThermoFisher Scientific | Cat# 12183020 |

| | | |
|---|---|---|
| Illumina polyA library preparation and NovaSeq PE sequencing | Genewiz/Azenta Life Sciences | N/A |
| **Deposited data** | | |
| RNA-seq | This work | GSE234019 |
| Single-cell RNA-seq | (Hung et al., 2020) | GSE120537 |
| Single-cell RNA-seq | (H. Li et al., 2022) | https://flycellatlas.org |
| **Experimental models: Organisms/strains** | | |
| *D. melanogaster*: *Su(H)GBE-lacZ; esg-Gal4, UAS-GFP, tubP-Gal80$^{ts}$/CyO; UAS-FLP, Act5C-FRT-CD2-FRT-Gal4/TM6C* (*esg$^{ts}$-FO* driver) | (Jiang et al., 2009) | N/A |
| *D. melanogaster*: *y, w; Su(H)GBE-Gal4, UAS-GFP, tubP-Gal80$^{ts}$/CyO; UAS-FLP, Act5C-FRT-CD2-FRT-Gal4/TM6B* (*NRE$^{ts}$-FO* driver) | (Zeng et al., 2010) | Derived from RRID:BDSC_83377 |
| *D. melanogaster*: *y, w; Su(H)GBE-Gal4/CyO; UAS-GFP, tubP-Gal80$^{ts}$/TM6B* (*NRE$^{ts}$* driver) | (Zeng et al., 2010) | Derived from RRID:BDSC_83377 |
| *D. melanogaster*: *UAS-da* | Sonsoles Campuzano (CSIC, Spain) | N/A |
| *D. melanogaster*: *UAS-sc* | Sonsoles Campuzano | N/A |
| *D. melanogaster*: *emc$^{EP3620}$* (*UAS-emc* gene trap) | Sonsoles Campuzano | N/A |
| *D. melanogaster*: *UAS-p35* | Sonsoles Campuzano | N/A |
| *D. melanogaster*: *UAS-da:da* | Sangbin Park (Stanford University, USA) | N/A |
| *D. melanogaster*: *UAS-N$^{ICD}$* | Alfonso Martínez Arias (UPF, Spain) | N/A |
| *D. melanogaster*: *UAS-N$^{RNAi}$* | (Presente et al., 2002) | N/A |
| *D. melanogaster*: *UAS-H* | Allison Bardin (Curie Institute, France) | N/A |
| *D. melanogaster*: *UAS-esg* | (Korzelius et al., 2014) | N/A |
| *D. melanogaster*: *UAS-Dcr-2* | Bloomington *Drosophila* Stock Center (BDSC) | RRID:BDSC_24646 |
| *D. melanogaster*: *y sc; UAS-da$^{RNAi}_{HMS01851}$* | BDSC | RRID:BDSC_38382 |
| *D. melanogaster*: *y v;; UAS-da$^{RNAi}_{JF02488}$* | BDSC | RRID:BDSC_29326 |
| *D. melanogaster*: *UAS-emc$^{RNAi}_{1007R-2}$* | National Institute of Genetics (Japan) | Stock# 1007R-2 |
| *D. melanogaster*: *UAS-emc$^{RNAi}_{KK108316}$* | Vienna *Drosophila* Resource Center | Stock# 100587 |
| *D. melanogaster*: *UAS-emc$^{RNAi}_{JF02300}$* | BDSC | RRID:BDSC_26738 |
| *D. melanogaster*: *UAS-H$^{RNAi}_{JF02624}$* | BDSC | RRID:BDSC_27315 |
| *D. melanogaster*: *UAS-esg$^{RNAi}_{HMS00025}$* | BDSC | RRID:BDSC_34063 |
| *D. melanogaster*: *emc$^{CPTI002740}$* | Kyoto *Drosophila* Stock Center (DGRC) | Stock# 115317 |
| *D. melanogaster*: *Myo1A-lacZ* | (Jiang et al., 2009) | RRID:BDSC_24646 |
| *D. melanogaster*: *da-GFP.FPTB* | BDSC | RRID:BDSC_55836 |
| *D. melanogaster*: *y w hs-Flp$_{1.22}$; Act5C-FRT-y$^{+}$-FRT-Gal4, UAS-lacZ$_{20b}$* | BDSC (modified) | RRID:BDSC_4410 |
| *D. melanogaster*: *y w hs-Flp$_{1.22}$ tub-Gal4 UAS-GFP; tub-Gal80 FRT40A / CyO* (MARCM FRT40A) | Allison Bardin | N/A |
| *D. melanogaster*: *y w hs-Flp$_{1.22}$ tub-Gal4 UAS-GFP; tub-Gal80 FRT80B / TM6B* (MARCM FRT80B) | Sonsoles Campuzano | N/A |
| *D. melanogaster*: *y w hs-Flp$_{1.22}$ tub-Gal4 UAS-GFP; tub-Gal80 FRT2A / TM6B* (MARCM FRT2A) | Sonsoles Campuzano | N/A |
| *D. melanogaster*: *w hs-Flp tub-Gal80 FRT19A; tub-Gal4 UAS-GFP / CyO* (MARCM FRT19A) | Shinya Yamamoto (Baylor College, USA) | N/A |
| *D. melanogaster*: *w; FRT40A* | BDSC | RRID:BDSC_1646 |
| *D. melanogaster*: *w;; FRT80B* | BDSC | RRID:BDSC_1620 |
| *D. melanogaster*: *y w;; FRT2A* | BDSC | RRID:BDSC_1997 |
| *D. melanogaster*: *y w FRT19A* | BDSC | RRID:BDSC_1709 |
| *D. melanogaster*: *w; Df(2L)da$^{10}$, FRT40A / In(2LR)Gla, Bc* | BDSC | RRID:BDSC_5531 |

| | | |
|---|---|---|
| *D. melanogaster*: *w;; emc$^{AP6}$ FRT80B / TM6B* | BDSC | RRID:BDSC_36544 |
| *D. melanogaster*: *w;; emc$^{1}$ FRT80B / TM2* | BDSC | RRID:BDSC_5532 |
| *D. melanogaster*: *emc$^{LL02590}$ FRT2A FRT82B / TM6C* | DGRC | Stock# 140642 |
| *D. melanogaster*: *Df(1)sc$^{B57}$ w FRT19A / FM7g* | Allison Bardin | N/A |
| Software and algorithms | | |
| Rstudio | (undefined author, n.d.) | http://www.rstudio.com |
| Illustrator CS6 | Adobe | N/A |
| Photoshop CS6 | Adobe | N/A |
| Designer 2 | Affinity | N/A |
| FIJI | (Schindelin et al., 2012) | https://fiji.sc |
| Other | | |
| Flygutseq | (Dutta et al., 2015) | https://flygutseq.buchonlab.com |
| Flybase | (Öztürk-Çolak et al., 2024) | https://flybase.org |

### RESOURCE AVAILABILITY

#### Lead contact

#### Materials availability

*Drosophila* strains generated in this study are available upon request. Requests for *Drosophila* strains should be directed to and will be fulfilled by the lead contact.

#### Data and code availability

- RNA-seq data have been deposited at GEO and are publicly available as of Jun 07, 2023. Accession number is in the key resources table. Microscopy data reported in this paper will be shared by the lead contact upon request.
- This paper does not report original code.
- Any additional information required to reanalyze the data reported in this work paper is available from the lead contact upon request.

### EXPERIMENTAL MODEL DETAILS

*Drosophila melanogaster* experimental subjects were adult mated females, aged for ~2 weeks. Subject females were housed with males at a ratio of 1:2 to 3:4 males to females, to allow mating, at a density of 5-9 flies/cm$^2$ of food surface and 2-3 flies/cm$^3$ of vial volume. Food was made from organic yellow maize flour (80 g/L), inactivated yeast powder (30 g/L), brewer's dextrose (80 g/L), agar (6.67 g/L), cooked at 95ºC before adding propionic acid (0.5%) and tegosept (0.005%). Vials were kept at 18ºC, 25ºC or 29ºC in a 12h:12h light/dark cycle. Breeding vials were flipped twice a week at 25ºC and once at 18ºC. Vials with adult experimental subjects were flipped every other day.

### METHOD DETAILS

#### Transgene and clonal induction

For experiments using Gal80$^{TS}$, adult flies were aged to gut maturity (4-7 days) at 18ºC, then transferred to 29 ºC. For induction of MARCM and flip-out clones, 4-7 days old flies were treated at 37ºC for 60 or 15 min, respectively. Flies were aged for 7 days after induction treatment before dissection, unless otherwise indicated (see Table S2). Fly strains are listed in the key resources table. All RNAi transgenes were co-expressed with *UAS-Dcr-2*.

#### Immunohistofluorescence

For antibody staining, adult guts were dissected in ice-cold PBS (maximum 10 min). Tissues were fixed in PBS-formaldehyde 4% (15 min at room temperature, RT), then in methanol (15 min RT). Methanol was washed off with three rinses in PBS-Triton X100 0.1% (PBT), followed by blocking and permeabilisation in PBT-BSA 2% (PBTB; three times, 15 min each). Tissues were incubated overnight at RT in primary antibody solution (diluted in PBTB to their final concentration; see key resources table). Primary antibody was washed off with three rinses and three 15-min incubations in PBT at RT. Tissue was incubated in PBT-secondary antibody solution (2 h at RT), then rinsed three times and incubated twice in PBT (15 min each); then once more in PBS. Tissue was equilibrated in mounting medium (4:1 glycerol:PBS with 4% w/v N-propyl-gallate) 4 h at RT or overnight at 4ºC. After mounting, 3D confocal imaging was performed in a Zeiss LSM 710 with an EC Plan-Neofluar 40X oil immersion objective (numerical aperture 1.3). Three positions along the anterior-posterior axis of each posterior midgut were acquired. In MARCM clone experiments, stacks were acquired from all clones found in each posterior midgut. Figures were assembled using FIJI and Adobe Photoshop/Illustrator CS6 or Affinity Designer 2.

**Cell counts**

For evaluating the proportion of cell types in GFP-labelled tissue, confocal stacks were maximum-intensity projected using FIJI and cells of the relevant types were counted manually with the Cell Counter plugin. Details of markers used can be found in **Figure S1D**. In the experiments co-expressing *sc* and *da:da*, the associated increase in proliferation generated large, highly densely populated cell clusters which could not be counted with single-cell precision. Therefore, for this genotype we estimated the proportion of each cell population in each field of view separately and then estimated the aggregated proportions.

**RNA-seq**

Flies bearing either *UAS-da*, *UAS-da*$^{RNAi}$ (TRiP.JF02092), *UAS-da:da* or *UAS-sc* as well as *esg-Gal4*, *UAS-GFP* and *tub-Gal80*$^{TS}$ were reared at 18ºC until 4-7 days old, transferred to 29ºC for 2 days and their midguts dissected, then processed as described in (Dutta et al., 2013). Libraries from three biological replicates per condition (except one condition, with two) were prepared in two batches and ~37 million reads (either 50 or 300bp long, for each batch respectively) were generated per library using Illumina technology. See key resources table for additional details.

**RNA-seq and scRNA-seq analysis**

Fastq read files, with adaptors pre-trimmed by the sequencing provider, were mapped to the release 6.28 of the *Drosophila melanogaster* genome using *STAR* and *bamtools* (Barnett et al., 2011; Dobin et al., 2013) and assigned to genes with *featureCounts* (Liao et al., 2014). Differential gene expression, gene set and DNA motif enrichment analyses were performed with the key R packages *DEseq2* (Love et al., 2014), *limma* (Ritchie et al., 2015), *fgsea* (Korotkevich et al., 2021) and *RcisTarget* (Aibar et al., 2017). We performed GSEA (Subramanian et al., 2005) against the Gene List Annotation for *Drosophila* (GLAD; (Hu et al., 2015), the Kyoto Encyclopedia of Genes and Genomes (KEGG; (Kanehisa et al., 2023) and gene lists from Dutta *et al*, (2015). scRNAseq data were obtained from GEO and https://flycellatlas.org. To correct for variability arising from technical and biological effects, we used the *IntegrateData* function in *Seurat* (Stuart et al., 2019; Hao et al., 2021). The *Slingshot* library (Street et al., 2018) was used for the trajectory analysis, which focused on ISC/EB cells as the initial state, pEC and EE cells, resulting in the identification of two distinct trajectories. The analysis code is in GitHub (https://github.com/jdenavascues/bHLH_code_midgut) and archived at Zenodo (doi:10.5281/zenodo.8116966).

**Quantification of Delta expression**

We took advantage of using simultaneously anti-Dl and anti-Pros mouse monoclonals, detected with the same anti-mouse secondary antibody, and of the robustness and reproducibility of the anti-Pros signal across conditions. This allows to use nuclear Pros staining outside the GFP-labelled MARCM clones as a normalisation reference, as the variation of Dl/Pros intensity ratio between samples is caused by the relative differences in Dl antigen. $Dl^+$ cells within clones and $Pros^+$ cells outside the clones were segmented so that we could take a value of fluorescence intensity per cell per marker, and normalised these values respect the average intensity of Pros per cell in that field of view.

***Segmentation and quantification in $Pros^+$ cells***

The positions of all cells were recorded in FIJI using CellCounter. A median filter was applied to the Pros/Dl channel to remove small features while preserving edges. A binary mask was created using Otsu thresholding (Otsu et al., 1979) of the filtered image. This mask captured most of the $Pros^+$ nuclei but missed some with lower expression. To segment these, we used the manually determined XY positions of the $Pros^+$ cells to add a 3-pixel diameter disk for each $Pros^+$ cell absent in the original mask. Fused nuclei were separated by marker-controlled watershed transformation (Meyer and Beucher, 1990). Pros expression for each nucleus was determined as the average pixel intensity value of the Pros signal channel in each segmented nucleus.

***Segmentation and quantification in $Dl^+$ cells***

Clones were detected by thresholding the GFP signal using the minimum cross-entropy method (C. H. Li and Lee, 1993). This missed a few cells, which were added to the mask using a similar approach to the $Pros^+$ nuclei, and the mask was consolidated by morphological filling and binary closing. Individual $Dl^+$ cells within the clone were identified by marker-controlled watershed segmentation, using the manually determined positions of the cells as markers. Dl expression for each cell was determined as the average pixel intensity of the Dl signal for each segmented object, normalised by the average Pros expression for that field of view.

### QUANTIFICATION AND STATISTICAL ANALYSIS

Statistical tests were performed in R. Change in cell type composition was assessed by binomial logistic regression for each individual cell type. In experiments with zero observations in one cell type, we used Firth's bias-reduced logistic regression (package *logistf*; (Heinze et al., 2023) to avoid the nonsensical results arising from the 'complete separation' of data (Albert and Anderson, 1984). All statistical tests and *p*-values of significance are specified in the corresponding figure legend; numbers of subjects are described either in the figure legend or **Tables S1-S3**. RNA-seq and scRNAseq analyses are described in detail in their methods detail section and the code repository indicated there.
