## Supplemental Figure Legends and Supplemental Tables 2 and 4 for "A bHLH interaction code controls bipotential differentiation and self-renewal in the *Drosophila* gut"

### SUPPLEMENTARY FIGURE LEGENDS

**Figure S1. Da monomers induce ISC/EB death by apoptosis.**

**A-B.** MARCM method: (A) the FRT-FLP system is used to induce recombination in G2, which during mitosis (M) can result in co-segregation of sister chromatid arms. (B) After induction, some ISCs generate sparse clones, allowing lineage tracing. **C.** The *esg$^{TS}$-FO* system allows simultaneous manipulation and tracing of ISCs/EBs. Top row: *esg-Gal4, UAS-GFP* is expressed specifically in ISCs/EBs. Bottom row: after induction, *Actin5C-Gal4* is reconstituted irreversibly in ISCs/EBs; they and their offspring will maintain GFP expression regardless of fate changes. **D.** Cell type markers used: ISCs are Dl$^{+}$ or Hdc$^{+}$/*NRE-lacZ*$^{—}$; EEs, Pros$^{+}$; preEEs, Dl$^{+}$/Pros$^{+}$; EBs, *NRE-lac*Z$^{+}$/Hdc$^{+}$; and ECs are *myoIA-lac*Z$^{+}$ and polyploid. **E.** Sequences used to generate *da$^{RNAi}$* and *da:da* transgenes; the *da$^{10}$* allele deletes the entire *da* locus.

**Figure S2. Da:Sc and Da:Da collaborate to induce proliferation.**

**A-B.** *sc* expression with *esg$^{TS}$-FO* (A) leads to formation of EEs and preEEs (A, empty asterisks and arrowheads, respectively) at the expense of EBs (A-B, solid arrowheads). Simultaneous knockdown of *da* (B) suppresses EE and preEE formation. Solid asterisks (-AB): extant EEs. **C.** Cell composition of GFP$^{+}$ tissue from A-B. **D-F.** Mitotic events (phospho-Histone 3, PH3) observed upon overexpression of *sc* (D), *sc* and *da* (E) or *sc* and tethered *da:da* (F) with *esg$^{TS}$-FO*. **G.** Quantification of mitosis in whole posterior midguts of wild-type and conditions in A-C. Numbers are 15, 10, 11 and 9 guts for wild-type, *sc*, *sc*+*da* and *sc*+*da:da*, respectively. *p*-values (Kruskall-Wallis/Dunn tests): ⁎ < 0.05; ⁎⁎ < 0.01; ⁎⁎⁎ < 0.001. DAI: days after induction. Scale bars: 20 μm.

**Figure S3. Da:Da and Da:Sc cooperate to induce the ISC transcriptional signature.**

**A.** PCA plot of RNAseq samples after log-normalisation and batch correction. PC1 segregates the overexpression of *sc* from the other conditions. PC2 is driven by the sequence *da$^{RNAi}$* < control < *da* < *da:da*. **B-C.** Log$_2$FC heatmap for the *E(spl)-Complex* genes (B) and 57 genes specific in expression or function for a midgut cell type (coloured as in **Figure 1I**) (C), for *sc*, *da$^{RNAi}$*, *da* and *da:da* RNAseq data. Non-HLH *E(spl)-C* genes are Broad Family Members (BFM). **D-E**. Normalised enrichment scores (NES) heatmaps for GLAD gene sets (D) or KEGG pathways (E). Coloured asterisks in heatmaps indicate significance as per each colonurmap. Column order follows hierarchical clustering.

**Figure S4. Sc confers ISC properties to EBs.**

**A.** The *$NRE^{TS}$-FO* system allows simultaneous manipulation and tracing of EBs. Top row: *NRE-Gal4, UAS-GFP* is expressed specifically in EBs. Bottom row: after induction, *Actin5C-Gal4* is reconstituted irreversibly in EBs; they and their offspring will maintain GFP expression regardless of fate changes. **B.** t-distributed stochastic neighbor embedding (t-SNE) representation of two integrated scRNAseq datasets from the adult intestine. All the annotated cell types are identified, and ISCs and EBs are distinguished from each other by the combined expression of *Dl*, *sna*, *polo* and *cnn* in ISCs, and *klu*, *E(spl)m3-HLH*, *E(spl)mα-BFM* and *E(spl)mβ-HLH* in EBs.

**Figure S5. *emc* is expressed in EBs/ECs and characterization of *$emc^{RNAi}$* transgenes.**

**A-H.** Gene expression by cell type, assessed with scRNAseq. *emc* (A) is expressed predominantly in EBs and posterior midgut ECs (pEC), with specificity similar to or higher than established EC markers *LManV*, *ι Try*, *β Try* and *myoIA* (B-E) and EB markers *E(spl)m3-HLH, klu* and *E(spl)mβ-HLH* (F-H). **I.** PCA plot with the transcriptional trajectories from ISCs/EBs into EEs or pECs derived from scRNAseq data. **J.** Molecular nature of alleles *$emc^{LL02590}$*, *$emc^{1}$*, *$emc^{AP6}$* and *$emc^{P5C}$*, and protein-trap *$emc^{CPTI002740}$*. **K-N.** Characterization of the efficiency of *$emc^{RNAi}$* transgenes *KK108316* and *JF02300* together (L) or *NIG-1007R* (M) expressed with *hh-Gal4*. Fold change in anti-Emc detection (N) is calculated as the ratio of nuclear staining in the anterior ($GFP^{-}$, left) wing compartment versus the affected posterior (GFP+, right) compartment, normalised across individual discs by the average anterior/posterior ratio in the wild-type contols (K). **O-R***.* Nota of flies bearing the *MS248* driver (O) and individual *$emc^{RNAi}$* transgenes, as indicated (P-R), showing increasingly strong *extra macrochaetae* phenotype.

**Figure S6. *emc* is required to prevent *$esg^{+}$* cell death and to repress Dl in EBs.**

**A-B.** Expressing *$emc^{RNAi}_{NIG}$* with *$esg^{TS}$-FO* leads to loss of ISCs/EBs (A); this is prevented by co-expression with *p35* (B). Arrowheads: *$esg^{+}$* cells. **C-D.** The *NRE-Gal4* (C) driver is expressed in EBs at high (solid arrowheads) and lower levels (empty arrowheads), but never in $Dl^{+}$ cells. In *NRE-Gal4, UAS-$emc^{RNAi}$* intestines (D), many EBs co-express Dl (empty arrowheads). DAI: days after induction. Scale bars: 20μm.

**Figure S7. *emc* and Notch work in parallel and there is no *da-esg* cross-regulation.**

**A-C.** Expressing $N^{RNAi}$ together with $da^{RNAi}$ (B) or *emc* (C) using $esg^{TS}$-*FO* leads to EC differentiation, abolishing the tumorous expansion of $Dl^{+}$ and $Pros^{+}$ cells with $N^{RNAi}$ alone (A). Solid arrowheads: polyploid ECs; empty arrowheads: $Dl^{-}$ diploid cell (EB or early EC). **D.** Cell composition of $GFP^{+}$ tissue from E-J and **Figure 5S**. *p*-values (binomial regression for individual cell types): ● < 0.05; ●● < 0.01; ●●● < 0.001. See **Tables S1-2** for statistical details. **E-J.** The co-expression of *emc* and *H* (F) or their double knockdown (I) with $esg^{TS}$-*FO* leads to phenotypes intermediate to the single-gene manipulations; co-expression of *H* and *emc* (F) increases EC differentiation (solid arrowheads) compared to *H* overexpression (E), but not as extensively as with *emc* (G); the simultaneous knock-down of *H* and *emc* (I) rescues the loss of ISCs (empty arrowheads) and EEs (asterisk) observed with $H^{RNAi}$ alone (H), but results in even further EC differentiation, and does not produce as many ISCs as with $emc^{RNAi}$ alone (J). **K.** Clonal expression with *Act5C-FO* of $N^{ICD}$, the cleavage-activated Notch reporter (blue) does not modify the expression of the GFP protein-trap line $emc^{CPTI002740}$. Solid arrowheads: ECs; empty arrowheads: EEs; asterisks: ISCs. **L-M.** Clonal expression with *Act5C-FO* of *esg* (L; blue) or *da:da* (M, green) does not modify the expression of the recombineering reporter *da:GFP* or the *lacZ* enhancer trap insertion $esg^{k00606}$, respectively. Solid arrowheads: *da* or *esg*-mis-expressing cells in L, M, respectively; empty arrowheads: non-mis-expressing cells. DAI: days after induction. Scale bars: 20µm. Panels A, E, G, H, J are reproduced from **Figures 2E, 7A, 5R, 7C, 5P**, respectively, to aid comparison.

### SUPPLEMENTARY TABLES

**Table S1. Binomial regression summary statistics (separate file – longer than 3 pages).**

**Table S2. Cell type proportions in the genotypes analysed (end of this file).**

**Table S3. RNAseq differentially expressed genes and gene counts (separate XLSX file).**

**Table S4. Cell type-exclusive markers (end of this file).**

**Table S5. List of enriched motifs in the genes upregulated or downregulated by overexpression of Da, Da:Da, Sc and $da^{RNAi}$, respectively (separate HTML file).** Motifs identified with RcisTarget using lists of genes differentially expressed at abs(log2FC) > 1.5.

### Table S2. Cell type proportions in the genotypes analysed

| | | Time | | | Tissue screened | | Cell Types (%) | | | | | | | |
|---|---|---|---|---|---|---|---|---|---|---|---|---|---|---|
| **Figure** | **Genotype** | **Days** | **since** (*) | **Total cells** | **No.** | **Tissue unit** (†) | **ISCs** | **EBs** | **EEs** | **ECs** | **PEEs** | **$Dl^+$ $NRE^+$** | **$Dl^-$ $NRE^-$** | **$Dl^+$ ECs** |
| **1A** | *FRT40A control* | 7 | induction | 443 | 12 | FoV | **12.0** | **39.7** | **6.5** | **41.8** | **0.0** | **0.0** | **0.0** | **0.0** |
| **1B** | *FRT40A da10* | 7 | induction | 495 | 54 | FoV | **2.0** | **4.6** | **1.0** | **92.3** | **0.0** | **0.0** | **0.0** | **0.0** |
| **1C** | *$esg^{TS}$-FO control* | 7 | induction | 1135 | 15 | FoV | **36.5** | **51.1** | **2.6** | **9.8** | **0.0** | **0.0** | **0.0** | **0.0** |
| **1D** | *$esg^{TS}$-FO > $da^{RNAi}{}_{JF}$* | 7 | induction | 803 | 15 | FoV | **0.0** | **0.2** | **1.6** | **98.1** | **0.0** | **0.0** | **0.0** | **0.0** |
| **1E** | *$esg^{TS}$-FO > $da^{RNAi}{}_{HMS}$* | 7 | induction | 1086 | 19 | FoV | **0.0** | **1.7** | **1.0** | **97.3** | **0.0** | **0.0** | **0.0** | **0.0** |
| **1F** | *$esg^{TS}$-FO > da* | 7 | induction | 549 | 14 | FoV | **23.9** | **51.7** | **24.2** | **0.2** | **0.0** | **0.0** | **0.0** | **0.0** |
| **1G** | *$esg^{TS}$-FO > da:da* | 7 | induction | 693 | 13 | FoV | **33.6** | **65.9** | **0.4** | **0.0** | **0.0** | **0.0** | **0.0** | **0.0** |
| **1H** | *$esg^{TS}$-FO > da:da + $da^{RNAi}{}_{JF}$* | 7 | induction | 1578 | 18 | FoV | **31.4** | **68.6** | **0.0** | **0.0** | **0.0** | **0.0** | **0.0** | **0.0** |
| **2A** | *$esg^{TS}$-FO > sc* | 3 | induction | 1241 | 6 | FoV | **3.9** | **2.7** | **33.8** | **1.4** | **58.3** | **0.0** | **0.0** | **0.0** |
| **2B** | *$esg^{TS}$-FO > sc + da* | 3 | induction | 2131 | 10 | FoV | **20.5** | **1.2** | **8.6** | **0.1** | **69.6** | **0.0** | **0.0** | **0.0** |
| **2C** | *$esg^{TS}$-FO > sc + da:da* (‡) | 3 | induction | 100 | 0 | FoV | **49.0** | **1.0** | **5.0** | **0.0** | **45.0** | **0.0** | **0.0** | **0.0** |
| **S2A** | *$esg^{TS}$-FO > sc* | 1 | induction | 553 | 11 | FoV | **36.3** | **37.4** | **1.8** | **6.5** | **17.9** | **0.0** | **0.0** | **0.0** |
| **S2B** | *$esg^{TS}$-FO > sc + $da^{RNAi}{}_{JF}$* | 1 | induction | 853 | 17 | FoV | **40.1** | **52.3** | **0.2** | **5.0** | **2.3** | **0.0** | **0.0** | **0.0** |
| **4A** | *$NRE^{TS}$-FO control* | 7 | induction | 1032 | 17 | FoV | **1.4** | **82.7** | **0.5** | **15.5** | **0.0** | **0.0** | **0.0** | **0.0** |
| **4B** | *$NRE^{TS}$-FO > da* | 2 | induction | 1563 | 20 | FoV | **14.4** | **84.6** | **0.1** | **0.5** | **0.4** | **0.0** | **0.0** | **0.0** |
| **4C** | *$NRE^{TS}$-FO > da:da* | 2 | induction | 460 | 4 | FoV | **6.5** | **92.8** | **0.7** | **0.0** | **0.0** | **0.0** | **0.0** | **0.0** |
| **4D** | *$NRE^{TS}$-FO > sc* | 7 | induction | 649 | 7 | FoV | **12.0** | **47.8** | **32.8** | **0.0** | **7.4** | **0.0** | **0.0** | **0.0** |
| **6I** | *$NRE^{TS}$-FO > $emc^{RNAi}{}_{NIG}$* | 7 | induction | 1628 | 16 | FoV | **76.4** | **23.5** | **0.0** | **0.1** | **0.0** | **0.0** | **0.0** | **0.0** |
| **5E** | *FRT2A control* | 7 | induction | 194 | 106 | clone | **34.5** | **45.4** | **4.6** | **15.5** | **0.0** | **0.0** | **0.0** | **0.0** |
| **5F** | *FRT2A $emc^{LL02590}$* | 7 | induction | 213 | 117 | clone | **60.6** | **32.4** | **3.8** | **3.3** | **0.0** | **0.0** | **0.0** | **0.0** |
| **5G** | *FRT80B $emc^{AP6}$* | 7 | induction | 208 | 110 | clone | **35.6** | **56.2** | **1.4** | **6.7** | **0.0** | **0.0** | **0.0** | **0.0** |
| **5H** | *FRT80B $emc^{1}$* | 7 | induction | 112 | 95 | clone | **44.6** | **46.4** | **8.0** | **0.9** | **0.0** | **0.0** | **0.0** | **0.0** |
| **S5M** | *FRT80B control* | 7 | induction | 310 | 117 | clone | **26.5** | **42.3** | **11.0** | **20.3** | **0.0** | **0.0** | **0.0** | **0.0** |
| **5J** | *$emc^{LL02590}$ / +* | 11-14 | emergence | 2964 | 10 | FoV | **16.6** | **17.9** | **5.6** | **59.8** | **0.0** | **0.0** | **0.0** | **0.0** |
| **5K** | *$emc^{LL02590}$ / $emc^{1}$* | 11-14 | emergence | 2488 | 12 | FoV | **20.0** | **20.5** | **8.7** | **48.8** | **3.0** | **0.0** | **0.0** | **1.9** |
| **5L** | *$emc^{LL02590}$ / $emc^{P5C}$* | 11-14 | emergence | 2582 | 10 | FoV | **22.2** | **21.9** | **6.2** | **40.0** | **1.0** | **0.0** | **0.0** | **9.7** |

| | | | | | | | | | | | | | | |
|---|---|---|---|---|---|---|---|---|---|---|---|---|---|---|
| **5O** | *esg$^{TS}$-FO > emc$^{RNAi}_{NIG}$ (III)* | 7 | induction | 104 | 22 | FoV | **83.7** | **5.8** | **9.6** | **1.0** | **0.0** | **0.0** | **0.0** | **0.0** |
| **5R** | *esg$^{TS}$-FO > emc$^{RNAi}_{NIG}$ (II)* | 7 | induction | 202 | 9 | FoV | **99.5** | **0.0** | **0.0** | **0.5** | **0.0** | **0.0** | **0.0** | **0.0** |
| **5P** | *esg$^{TS}$-FO > emc$^{RNAi}_{KK+JF}$* | 7 | induction | 832 | 15 | FoV | **41.2** | **11.3** | **0.8** | **1.7** | **0.0** | **45.0** | **0.0** | **0.0** |
| **5Q** | *esg$^{TS}$-FO > emc* | 7 | induction | 812 | 10 | FoV | **0.0** | **0.2** | **0.4** | **99.4** | **0.0** | **0.0** | **0.0** | **0.0** |
| **6C** | *FRT2A emc$^{LL02590}$ + da$^{RNAi}_{JF}$* | 7 | induction | 412 | 35 | FoV | **3.9** | **12.1** | **0.7** | **83.3** | **0.0** | **0.0** | **0.0** | **0.0** |
| **6C** | *FRT40A da$^{10}$ + emc$^{RNAi}_{NIG}$* | 7 | induction | 137 | 23 | FoV | **0.7** | **3.6** | **1.5** | **94.2** | **0.0** | **0.0** | **0.0** | **0.0** |
| **7A** | *esg$^{TS}$-FO > H* | 7 | induction | 1810 | 21 | FoV | **77.5** | **17.0** | **3.7** | **1.8** | **0.0** | **0.0** | **0.0** | **0.0** |
| **7B** | *esg$^{TS}$-FO > H + da$^{RNAi}_{JF}$* | 7 | induction | 1510 | 21 | FoV | **18.2** | **0.1** | **1.2** | **78.2** | **0.0** | **0.0** | **2.3** | **0.0** |
| **7C** | *esg$^{TS}$-FO > H$^{RNAi}_{HMS}$* | 7 | induction | 984 | 15 | FoV | **0.0** | **55.7** | **0.5** | **26.1** | **0.0** | **17.7** | **0.0** | **0.0** |
| **7D** | *esg$^{TS}$-FO > H$^{RNAi}_{HMS}$ + da:da* | 7 | induction | 613 | 8 | FoV | **0.0** | **99.2** | **0.8** | **0.0** | **0.0** | **0.0** | **0.0** | **0.0** |
| **7E** | *esg$^{TS}$-FO > esg$^{RNAi}_{HMS}$* | 7 | induction | 628 | 18 | FoV | **0.2** | **0.3** | **7.5** | **92.0** | **0.0** | **0.0** | **0.0** | **0.0** |
| **7F** | *esg$^{TS}$-FO > esg$^{RNAi}_{HMS}$ + da:da* | 7 | induction | 939 | 19 | FoV | **41.4** | **56.8** | **1.8** | **0.0** | **0.0** | **0.0** | **0.0** | **0.0** |
| **7G** | *esg$^{TS}$-FO > esg* | 7 | induction | 850 | 9 | FoV | **34.1** | **65.8** | **0.1** | **0.0** | **0.0** | **0.0** | **0.0** | **0.0** |
| **7H** | *esg$^{TS}$-FO > esg + da$^{RNAi}_{JF}$* | 7 | induction | 1236 | 12 | FoV | **61.7** | **38.3** | **0.0** | **0.0** | **0.0** | **0.0** | **0.0** | **0.0** |
| **S7F** | *esg$^{TS}$-FO > H + emc* | 7 | induction | 1292 | 27 | FoV | **44.2** | **19.2** | **0.0** | **36.6** | **0.0** | **0.0** | **0.0** | **0.0** |
| **S7I** | *esg$^{TS}$-FO > H$^{RNAi}_{HMS}$ + emc$^{RNAi}_{NIG}$* | 7 | induction | 126 | 16 | FoV | **31.7** | **11.9** | **5.6** | **46.0** | **0.0** | **4.8** | **0.0** | **0.0** |

* Induction consists of a temperature switch for Flip-Out genotypes and a heat shock of 60 min at 37ºC for MARCM genotypes.

† FoV: field of view (40X objective).

‡ Estimated (see methods).

**Table S4. Cell type-exclusive markers.**

| Cell type | Gene symbol | Reference (function) | Reference (expression) | FCA[§] marker |
|---|---|---|---|---|
| EB | *klu* | (Korzelius et al., 2019) | (Korzelius et al., 2019; Hung et al., 2020) | TRUE |
| EB | *E(spl)mbeta-HLH* | (Lu and Z. Li, 2015) | (Lu and Z. Li, 2015) | TRUE |
| EB | *CG42747* | NA | (Li et al., 2022) | TRUE |
| EB | *Dtg* | NA | (Li et al., 2022) | TRUE |
| EB | *CG14457* | NA | (Li et al., 2022) | TRUE |
| EB | *CG30090* | NA | (Li et al., 2022) | TRUE |
| EB | *CG34236* | NA | (Li et al., 2022) | TRUE |
| EB | *lncRNA:CR43270* | NA | (Li et al., 2022) | TRUE |
| EB | *CDase* | NA | (Li et al., 2022) | TRUE |
| EB | *E(spl)m3-HLH* | NA | (Li et al., 2022) | FALSE |
| EB | *Lrch* | NA | (Li et al., 2022) | FALSE |
| pEC | *alphaTry* | NA | (Hung et al., 2020) | TRUE |
| pEC | *Amy-d* | NA | (Hung et al., 2020) | TRUE |
| pEC | *Amy-p* | NA | (Hung et al., 2020) | TRUE |
| pEC | *betaTry* | NA | (Hung et al., 2020) | TRUE |
| pEC | *ck* | NA | (Jia Chen et al., 2018) | FALSE |
| pEC | *LManV* | NA | (Hung et al., 2020) | FALSE |
| pEC | *LManVI* | NA | (Hung et al., 2020) | TRUE |
| pEC | *Myo31DF* | NA | (Hung et al., 2020; Jiang et al., 2009) | TRUE |
| pEC | *nub* | (Dantoft et al., 2013) | (Hung et al., 2020; Lee et al., 2009) | TRUE |
| pEC | *Ssk* | (Hsi-Ju Chen et al., 2020) | (Hung et al., 2020) | TRUE |
| pEC | *iotaTry* | NA | (Hung et al., 2020) | TRUE |
| pEC | *CG14219* | NA | (Li et al., 2022) | TRUE |
| pEC | *CG13492* | NA | (Li et al., 2022) | TRUE |
| pEC | *CG10911* | NA | (Li et al., 2022) | TRUE |
| pEC | *Mal-A4* | NA | (Li et al., 2022) | TRUE |
| pEC | *Acbp3* | NA | (Li et al., 2022) | TRUE |
| pEC | *Burs* | NA | (Li et al., 2022) | TRUE |
| pEC | *Mal-A3* | NA | (Li et al., 2022) | TRUE |
| EE | *AstA* | NA | (Ohlstein and Spradling, 2006; Hung et al., 2020) | TRUE |
| EE | *Mip* | NA | (Beehler-Evans and Micchelli, 2015) | FALSE |
| EE | *AstC* | NA | (Ohlstein and Spradling, 2006; Hung et al., 2020) | TRUE |
| EE | *Dh31* | NA | (Beehler-Evans and Micchelli, 2015; Chen et al., 2016) | FALSE |
| EE | *dimm* | (Beebe et al., 2015) | (Hung et al., 2020; Beebe et al., 2015) | TRUE |
| EE | *mirr* | NA | (Guo et al., 2022) | FALSE |
| EE | *NPF* | NA | (Chen et al., 2016; Hung et al., 2020) | TRUE |
| EE | *phyl* | (Yin and Xi, 2018) | NA | FALSE |
| EE | *Poxn* | NA | (Dutta et al., 2015) | FALSE |
| EE | *pros* | (Biteau and Jasper, 2014) | (Ohlstein and Spradling, 2006; Micchelli and Perrimon, 2006) | FALSE |

| | | | | |
|---|---|---|---|---|
| EE | *Ptx1* | NA | (Guo et al., 2022) | FALSE |
| EE | *sina* | (Yin and Xi, 2018) | NA | FALSE |
| EE | *tap* | (Hartenstein et al., 2017) | (Hartenstein et al., 2017) | FALSE |
| EE | *Tk* | NA | (Ohlstein and Spradling, 2006; Hung et al., 2020) | TRUE |
| EE | *CG34386* | NA | (Li et al., 2022) | TRUE |
| EE | *TrpA1* | NA | (Li et al., 2022) | TRUE |
| EE | *Orcokinin* | NA | (Chen et al., 2016) | TRUE |
| EE | *CG7191* | NA | (Li et al., 2022) | TRUE |
| EE | *CngA* | NA | (Li et al., 2022) | TRUE |
| EE | *CCHa1* | NA | (Li et al., 2022) | TRUE |
| EE | *Sytbeta* | NA | (Li et al., 2022) | TRUE |
| ISC | *Dl* | (Ohlstein and Spradling, 2007) | (Ohlstein and Spradling, 2007) | TRUE |
| ISC | *mira* | NA | (Bardin et al., 2010) | FALSE |
| ISC | *polo* | NA | (Amcheslavsky et al., 2011, 2014) | FALSE |
| ISC | *spdo* | NA | (Perdigoto et al., 2011) | FALSE |
| ISC | *zfh2* | (Rojas Villa et al., 2019) | (Doupé et al., 2018; Dutta et al., 2015) | TRUE |
| ISC | *mew* | NA | (Li et al., 2022) | TRUE |
| ISC | *Trim9* | NA | (Li et al., 2022) | TRUE |

§ FCA: Fly Cell Atlas
