## Supplemental Table 1 for "A bHLH interaction code controls bipotential differentiation and self-renewal in the *Drosophila* gut"

### Table S1. Binomial regression summary statistics

| Figures | Allele/transgene | Reference genotype | Time (*) | Cell type tested | Effect | Odds Ratio | *p*-value (†) |
|---|---|---|---|---|---|---|---|
| **1I** | *$FRT_{40A}$ $da^{10}$* | *$FRT_{40A}$ control* | 7 DAI | **ISC** | ↓↓↓ | 0.15 | 8e-08 |
| | | | | **EB** | ↓↓↓ | 0.07 | 1.2e-28 |
| | | | | **EE** | ↓↓↓ | 0.15 | 8.1e-05 |
| | | | | **EC** | ↑↑↑ | 17 | 1.1e-47 |
| | *$esg^{TS}$-FO > $da^{RNAi}{}_{JF}$* | *$esg^{TS}$-FO control* | 7 DAI | **ISC** | ↓↓↓ (f) | 1.1e-03 | 0 |
| | | | | **EB** | ↓↓↓ | 2.4e-03 | 1.9e-17 |
| | | | | **EE** | — | 0.61 | 0.14 |
| | | | | **EC** | ↑↑↑ | 490 | 9.7e-109 |
| | *$esg^{TS}$-FO > $da^{RNAi}{}_{HMS}$* | *$esg^{TS}$-FO control* | 7 DAI | **ISC** | ↓↓↓ (f) | 8e-04 | 0 |
| | | | | **EB** | ↓↓↓ | 0.016 | 1e-63 |
| | | | | **EE** | ↓↓ | 0.38 | 6e-03 |
| | | | | **EC** | ↑↑↑ | 340 | 4.2e-164 |
| | *$esg^{TS}$-FO > da* | *$esg^{TS}$-FO control* | 7 DAI | **ISC** | ↓↓↓ | 0.55 | 2.6e-07 |
| | | | | **EB** | — | 1 | 0.81 |
| | | | | **EE** | ↑↑↑ | 12 | 8.4e-32 |
| | | | | **EC** | ↓↓↓ | 0.017 | 4.8e-05 |
| | *$esg^{TS}$-FO > da:da* | *$esg^{TS}$-FO control* | 7 DAI | **ISC** | — | 0.89 | 0.22 |
| | | | | **EB** | ↑↑↑ | 1.9 | 6.3e-10 |
| | | | | **EE** | ↓↓ | 0.16 | 2.6e-03 |
| | | | | **EC** | ↓↓↓ (f) | 6.7e-03 | 0 |
| | *$esg^{TS}$-FO > da:da + $da^{RNAi}{}_{JF}$* | *$esg^{TS}$-FO control* | 7 DAI | **ISC** | ↓↓ | 0.8 | 6.1e-03 |
| | | | | **EB** | ↑↑↑ | 2.1 | 5.6e-20 |
| | | | | **EE** | ↓↓↓ (f) | 0.012 | 2.5e-12 |
| | | | | **EC** | ↓↓↓ (f) | 3e-03 | 0 |

| | | | | | | | |
|---|---|---|---|---|---|---|---|
| **2D** | *esg$^{TS}$-FO > sc + da* | *esg$^{TS}$-FO > sc* | 3 DAI | **ISC** | ↑↑↑ | 6.3 | 2.8e-32 |
| | | | | **EB** | ↓↓ | 0.44 | 1.8e-03 |
| | | | | **EE** | ↓↓↓ | 0.18 | 6.9e-67 |
| | | | | **EC** | ↓↓↓ | 0.1 | 2.7e-04 |
| | | | | **PEE** | ↑↑↑ | 1.6 | 3e-11 |
| **S2C** | *esg$^{TS}$-FO > sc + da$^{RNAi}_{JF}$* | *esg$^{TS}$-FO > sc* | 1 DAI | **ISC** | — | 1.2 | 0.16 |
| | | | | **EB** | ↑↑↑ | 1.8 | 5.6e-08 |
| | | | | **EE** | ↓↓ | 0.13 | 8e-03 |
| | | | | **EC** | — | 0.76 | 0.24 |
| | | | | **PEE** | ↓↓↓ | 0.11 | 2e-18 |
| **4E** | *NRE$^{TS}$-FO > da* | *NRE$^{TS}$-FO control* | 7 DAI | **ISC** | ↑↑↑ | 12 | 2.5e-19 |
| | | | | **EB** | — | 1.2 | 0.19 |
| | | | | **EE** | — | 0.13 | 0.064 |
| | | | | **EC** | ↓↓↓ | 0.028 | 1.1e-22 |
| | | | | **PEE** | ↑ *(f)* | 10 | 0.028 |
| | *NRE$^{TS}$-FO > da:da* | *NRE$^{TS}$-FO control* | 7 DAI | **ISC** | ↑↑↑ | 5.1 | 7.8e-07 |
| | | | | **EB** | ↑↑↑ | 2.7 | 4.8e-07 |
| | | | | **EE** | — | 1.4 | 0.68 |
| | | | | **EC** | ↓↓↓ *(f)* | 5.9e-03 | 0 |
| | *NRE$^{TS}$-FO > sc* | *NRE$^{TS}$-FO control* | 7 DAI | **ISC** | ↑↑↑ | 9.9 | 7e-15 |
| | | | | **EB** | ↓↓↓ | 0.19 | 9.6e-48 |
| | | | | **EE** | ↑↑↑ | 100 | 5.2e-24 |
| | | | | **EC** | ↓↓↓ *(f)* | 4.2e-03 | 0 |
| | | | | **PEE** | ↑↑↑ *(f)* | 170 | 0 |
| **5I** | *FRT$_{2A}$ emc$^{LL02590}$* | *FRT$_{2A}$ control* | 7 DAI | **ISC** | ↑↑↑ | 2.9 | 2.2e-07 |
| | | | | **EB** | ↓↓ | 0.58 | 7.5e-03 |
| | | | | **EE** | — | 0.8 | 0.66 |
| | | | | **EC** | ↓↓↓ | 0.19 | 1e-04 |

| | | | | | | | |
|---|---|---|---|---|---|---|---|
| | *FRT$_{80B}$ emc$^{AP6}$* | *FRT$_{80B}$ control* | 7 DAI | **ISC** | ↑ | 1.5 | 0.027 |
| | | | | **EB** | ↑↑ | 1.8 | 1.8e-03 |
| | | | | **EE** | ↓↓↓ | 0.12 | 4.7e-04 |
| | | | | **EC** | ↓↓↓ | 0.28 | 4.8e-05 |
| | *FRT$_{80B}$ emc$^{1}$* | *FRT$_{80B}$ control* | 7 DAI | **ISC** | ↑↑↑ | 2.2 | 4.4e-04 |
| | | | | **EB** | — | 1.2 | 0.45 |
| | | | | **EE** | — | 0.71 | 0.38 |
| | | | | **EC** | ↓↓↓ | 0.035 | 9.8e-04 |
| **5M** | *emc$^{LL02590}$ / emc$^{1}$* | *emc$^{LL02590}$ / +* | 11-14 DAE | **ISC** | ↑↑ | 1.3 | 1.4e-03 |
| | | | | **EB** | ↑ | 1.2 | 0.017 |
| | | | | **EE** | ↑↑↑ | 1.6 | 8.3e-06 |
| | | | | **EC** | ↓↓↓ | 0.64 | 4.4e-16 |
| | | | | **PEE** | — (f) | 8.4 | 0.079 |
| | | | | **Dl$^{+}$ EC** | ↑↑↑ | 58 | 5.7e-05 |
| | *emc$^{LL02590}$ / emc$^{P5C}$* | *emc$^{LL02590}$ / +* | 11-14 DAE | **ISC** | ↑↑↑ | 1.4 | 1.7e-07 |
| | | | | **EB** | ↑↑↑ | 1.3 | 2.2e-04 |
| | | | | **EE** | — | 1.1 | 0.35 |
| | | | | **EC** | ↓↓↓ | 0.45 | 1.4e-48 |
| | | | | **PEE** | — (f) | 3.5 | 0.41 |
| | | | | **Dl$^{+}$ EC** | ↑↑↑ | 320 | 8.8e-09 |
| **5R** | *esg$^{TS}$-FO > emc$^{RNAi}$$_{NIG}$ (III)* | *esg$^{TS}$-FO control* | 7 DAI | **ISC** | ↑↑↑ | 8.9 | 9.4e-16 |
| | | | | **EB** | ↓↓↓ | 0.059 | 2.4e-11 |
| | | | | **EE** | ↑↑↑ | 3.9 | 3.3e-04 |
| | | | | **EC** | ↓ | 0.09 | 0.017 |
| **5R** | *esg$^{TS}$-FO > emc$^{RNAi}$$_{NIG}$ (II)* | *esg$^{TS}$-FO control* | 7 DAI | **ISC** | ↑↑↑ | 350 | 5.3e-09 |
| | | | | **EB** | ↓↓↓ (f) | 2.4e-03 | 0 |
| | | | | **EE** | ↓↓ (f) | 0.09 | 8.6e-03 |
| | | | | **EC** | ↓↓ | 0.046 | 2.2e-03 |

| | | | | | | | |
|---|---|---|---|---|---|---|---|
| | $esg^{TS}$-FO > $emc^{RNAi}_{KK+JF}$ | $esg^{TS}$-FO control | 7 DAI | **ISC** | ↑ | 1.2 | 0.033 |
| | | | | **EB** | ↓↓↓ | 0.12 | 4.9e-64 |
| | | | | **EE** | ↓↓ | 0.31 | 5.9e-03 |
| | | | | **EC** | ↓↓↓ | 0.16 | 1.3e-10 |
| | | | | **$Dl^{+}NRE^{+}$** | ↑↑↑ (f) | 1850 | 0 |
| | $esg^{TS}$-FO > emc | $esg^{TS}$-FO control | 7 DAI | **ISC** | ↓↓↓ (f) | 1.1e-03 | 0 |
| | | | | **EB** | ↓↓↓ | 2.4e-3 | 1.7e-17 |
| | | | | **EE** | ↓↓ | 0.14 | 1e-03 |
| | | | | **EC** | ↑↑↑ | 1500 | 6.3e-57 |
| **6E** | $FRT_{40A}$ $da^{10}$ + $emc^{RNAi}_{NIG}$ | $FRT_{40A}$ control | 7 DAI | **ISC** | ↓↓ | 0.054 | 4e-03 |
| | | | | **EB** | ↓↓↓ | 0.058 | 8.5e-10 |
| | | | | **EE** | ↓ | 0.21 | 0.035 |
| | | | | **EC** | ↑↑↑ | 23 | 1.5e-16 |
| | $FRT_{40A}$ $da^{10}$ + $emc^{RNAi}_{NIG}$ | $FRT_{40A}$ $da^{10}$ | 7 DAI | **ISC** | — | 0.36 | 0.33 |
| | | | | **EB** | — | 0.78 | 0.62 |
| | | | | **EE** | — | 1.6 | 0.66 |
| | | | | **EC** | — | 1.3 | 0.47 |
| | $FRT_{2A}$ $emc^{LL02590}$ + $da^{RNAi}_{JF}$ | $FRT_{2A}$ control | 7 DAI | **ISC** | ↓↓↓ | 0.077 | 4.3e-18 |
| | | | | **EB** | ↓↓↓ | 0.17 | 8.4e-18 |
| | | | | **EE** | ↓↓ | 0.15 | 4.9e-03 |
| | | | | **EC** | ↑↑↑ | 27 | 1.2e-43 |
| | $FRT_{2A}$ $emc^{LL02590}$ + $da^{RNAi}_{JF}$ | $FRT_{2A}$ $emc^{LL02590}$ | 7 DAI | **ISC** | ↓↓↓ | 0.026 | 7.4e-36 |
| | | | | **EB** | ↓↓↓ | 0.29 | 3.3e-09 |
| | | | | **EE** | ↓ | 0.19 | 0.014 |
| | | | | **EC** | ↑↑↑ | 150 | 1.3e-34 |
| **6J** | $NRE^{TS}$-FO > $emc^{RNAi}_{NIG}$ | $NRE^{TS}$-FO control | 7 DAI | **ISC** | ↑↑↑ | 240 | 1.8e-87 |
| | | | | **EB** | ↓↓↓ | 0.065 | 1.5e-162 |
| | | | | **EE** | ↓↓ (f) | 0.058 | 6.2e-03 |
| | | | | **EC** | ↓↓↓ | 6.7e-03 | 2.2e-12 |

| | | | | | | | |
|---|---|---|---|---|---|---|---|
| **7I** | *esg<sup></sup>*$esg^{TS}$-FO > H | $esg^{TS}$-FO > H + $da^{RNAi}_{JF}$ | 7 DAI | **ISC** | ↑↑↑ | 16 | 2.5e-216 |
| | | | | **EB** | ↑↑↑ | 310 | 1e-08 |
| | | | | **EE** | ↑↑↑ | 3.2 | 1.5e-05 |
| | | | | **EC** | ↓↓↓ | 5e-03 | 6.8e-173 |
| | | | | **Dl⁻NRE⁻** | ↓↓↓ (f) | 0.012 | 6.4e-13 |
| | $esg^{TS}$-FO > $da^{RNAi}_{JF}$ | $esg^{TS}$-FO > H + $da^{RNAi}_{JF}$ | 7 DAI | **ISC** | ↓↓↓ (f) | 2.8e-03 | 0 |
| | | | | **EB** | — | 3.8 | 0.28 |
| | | | | **EE** | — | 1.4 | 0.4 |
| | | | | **EC** | ↑↑↑ | 15 | 1.3e-23 |
| | | | | **Dl⁻NRE⁻** | ↓↓↓ (f) | 0.026 | 2.8e-07 |
| | $esg^{TS}$-FO > $H^{RNAi}_{HMS}$ | $esg^{TS}$-FO > $H^{RNAi}_{HMS}$ + da:da | 7 DAI | **EB** | ↓↓↓ | 0.01 | 6.8e-24 |
| | | | | **EE** | — | 0.62 | 0.45 |
| | | | | **EC** | ↑↑↑ (f) | 430 | 0 |
| | | | | **Dl⁺NRE⁺** | ↑↑↑ (f) | 260 | 0 |
| | $esg^{TS}$-FO > da:da | $esg^{TS}$-FO > $H^{RNAi}_{HMS}$ + da:da | 7 DAI | **ISC** | ↑↑↑ (f) | 620 | 0 |
| | | | | **EB** | ↓↓↓ | 0.016 | 1.1e-19 |
| | | | | **EE** | — | 0.53 | 0.38 |
| **7J** | $esg^{TS}$-FO > $esg^{RNAi}_{HMS}$ | $esg^{TS}$-FO > $esg^{RNAi}_{HMS}$ + da:da | 7 DAI | **ISC** | ↓↓↓ | 2.3e-03 | 1.2e-09 |
| | | | | **EB** | ↓↓↓ | 2.4e-03 | 2.6e-17 |
| | | | | **EE** | ↑↑↑ | 4.4 | 2.8e-07 |
| | | | | **EC** | ↑↑↑ (f) | 21500 | 0 |
| | $esg^{TS}$-FO > da:da | $esg^{TS}$-FO > $esg^{RNAi}_{HMS}$ + da:da | 7 DAI | **ISC** | ↓↓ | 0.72 | 1.4e-03 |
| | | | | **EB** | ↑↑↑ | 1.5 | 1.8e-04 |
| | | | | **EE** | ↓ | 0.24 | 0.021 |
| | $esg^{TS}$-FO > esg | $esg^{TS}$-FO > esg + $da^{RNAi}_{JF}$ | 7 DAI | **ISC** | ↓↓↓ | 0.32 | 4.2e-34 |
| | | | | **EB** | ↑↑↑ | 3.1 | 7.7e-34 |
| | | | | **EE** | — (f) | 4.4 | 0.33 |

| | | | | | | | |
|---|---|---|---|---|---|---|---|
| | *esg$^{TS}$-FO > da$^{RNAi}$$_{JF}$* | *esg$^{TS}$-FO > esg + da$^{RNAi}$$_{JF}$* | 7 DAI | **ISC** | ↓↓↓ $^{(f)}$ | 3.9e-04 | 0 |
| | | | | **EB** | ↓↓↓ | 4e-03 | 7.9e-15 |
| | | | | **EE** | ↑↑↑ $^{(f)}$ | 42 | 3.6e-06 |
| | | | | **EC** | ↑↑↑ $^{(f)}$ | 126000 | 0 |
| **S7D** | *esg$^{TS}$-FO > H* | *esg$^{TS}$-FO > H + emc* | 7 DAI | **ISC** | ↑↑↑ | 4.4 | 1.5e-76 |
| | | | | **EB** | — | 0.86 | 0.12 |
| | | | | **EE** | ↑↑↑ $^{(f)}$ | 100 | 1.1e-16 |
| | | | | **EC** | ↓↓↓ | 0.031 | 1.9e-76 |
| | *esg$^{TS}$-FO > emc* | *esg$^{TS}$-FO > H + emc* | 7 DAI | **ISC** | ↓↓↓ $^{(f)}$ | 7.8e-04 | 0 |
| | | | | **EB** | ↓↓↓ | 0.01 | 1.4e-10 |
| | | | | **EE** | ↑ $^{(f)}$ | 11 | 0.042 |
| | | | | **EC** | ↑↑↑ | 280 | 1.3e-35 |
| | *esg$^{TS}$-FO > H$^{RNAi}$$_{HMS}$* | *esg$^{TS}$-FO > H$^{RNAi}$$_{HMS}$ + emc$^{RNAi}$$_{NIG}$* | 7 DAI | **ISC** | ↓↓↓ $^{(f)}$ | 1.1e-03 | 0 |
| | | | | **EB** | ↑↑↑ | 9.3 | 2.9e-15 |
| | | | | **EE** | ↓↓↓ | 0.087 | 3.8e-05 |
| | | | | **EC** | ↓↓↓ | 0.41 | 5e-06 |
| | | | | **Dl$^{+}$NRE$^{+}$** | ↑↑↑ | 4.3 | 6.3e-04 |
| | *esg$^{TS}$-FO > emc$^{RNAi}$$_{NIG}$ (III)* | *esg$^{TS}$-FO > H$^{RNAi}$$_{HMS}$ + emc$^{RNAi}$$_{NIG}$* | 7 DAI | **ISC** | ↑↑↑ | 11 | 2.2e-13 |
| | | | | **EB** | — | 0.45 | 0.12 |
| | | | | **EE** | — | 1.8 | 0.25 |
| | | | | **EC** | ↓↓↓ | 0.011 | 1.2e-05 |
| | | | | **Dl$^{+}$NRE$^{+}$** | ↓ $^{(f)}$ | 0.089 | 0.023 |

* DAI, *d*ays *a*fter transgene or clone *i*nduction. DAE, *d*ays *a*fter *e*mergence.

† ↑, ↑↑, or ↑↑↑ indicate $p < 0.05$, $p < 0.01$ or $p < 0.001$, respectively, for increased cell type frequency; ↓ is used instead of ↑ to indicate significance for decreased cell type frequency.

$^{f}$ Cases where Firth's bias reduced logistic regression was used. See Methods.
